# *Mycobacterium tuberculosis* inactivates host oxysterols to impair macrophage cholesterol homeostasis

**DOI:** 10.64898/2026.10.04.756586

**Authors:** Andrew T. Roth, Mohsen Ali Asgari, Pallavi Chandra, Jalalah Muhammad, Amy Lopez, Sourav Bhattacharya, Isabelle A. Williams, Brian T. Edelson, Charles W. Goss, Dmitri I. Kotov, Mingxing Qian, Douglas F. Covey, Xuntian Jiang, William J. Griffiths, Yuqin Wang, Jennifer A. Philips

## Abstract

Host oxysterols coordinate macrophage cholesterol homeostasis and antimicrobial defense, but whether pathogens directly target oxysterol signaling is unknown. Here, we identify pathogen-mediated enzymatic oxysterol inactivation as a mechanism by which *Mycobacterium tuberculosis* (Mtb) subverts host cholesterol metabolism. Mtb oxidizes the endogenous oxysterols 27-hydroxycholesterol (27-HC) and 3β-hydroxycholest-5-enoic acid (3β-HCA) into 3-oxo-Δ^4^ metabolites. Sputum lipidomics across TB cohorts on three continents revealed an active-disease signature characterized by 27-HC depletion and accumulation of 27-hydroxycholest-4-en-3-one (27-HCO) and 3-oxocholest-4-enoic acid (3O-CA), which normalized with treatment. In infected human macrophages, Mtb 3β-hydroxysteroid dehydrogenase (3β-Hsd) generated 27-HCO. Unlike their parent oxysterols, 27-HCO and 3O-CA did not activate liver X receptor (LXR), and 3O-CA failed to suppress sterol regulatory element-binding protein 2 (SREBP2) target genes. Accordingly, 3β-Hsd-deficient Mtb did not induce macrophage cholesterol retention, a phenotype restored by LXR antagonism. Thus, Mtb enzymatically inactivates host oxysterols, disrupting macrophage cholesterol homeostasis and generating treatment-responsive metabolic signatures in human TB.

## Introduction

Tuberculosis (TB), an infection caused by *Mycobacterium tuberculosis* (Mtb), remains the leading infectious disease killer worldwide, claiming more than 1 million lives each year.^1^ Mtb establishes infection in alveolar macrophages, cells that normally clear inhaled microbes, cellular debris, and excess pulmonary surfactant.^2,3^ To prevent lipid overload, lung macrophages efflux excess cholesterol to extracellular lipid acceptors and metabolize cholesterol to more soluble 27-oxygenated sterols that can enter the circulation for hepatic metabolism and excretion.^3–6^ Mtb remodels macrophage lipid homeostasis, promoting the formation of lipid-overloaded “foamy” macrophages that accumulate triacylglycerols along with cholesteryl esters.^7^ These cells are thought to provide nutrient-rich niche that supports Mtb persistence, and in some cases exhibit impaired antimicrobial functions. Mtb can also import and catabolize host cholesterol to support bacterial metabolism during chronic infection.^8–18^ However, the mechanisms by which Mtb disrupts host cholesterol homeostasis are incompletely understood.

Oxysterols are oxygenated cholesterol derivatives that regulate lipid handling, inflammatory responses, and host defense.^19–22^ In the lungs, 27-hydroxycholesterol (27-HC), also known as (25R)26-hydroxycholesterol, is a major regulator of cholesterol homeostasis. 27-HC is generated by sterol 27-hydroxylase (CYP27A1), which can further oxidize 27-HC to 3β-hydroxycholest-5-enoic acid (3β-HCA), a precursor of C_24_ bile acids.^22,23^ Oxysterols, including 27-HC, impair the sterol regulatory element-binding protein 2 (SREBP2) pathway,^24^ which promotes cholesterol uptake and synthesis,^20,25,26^ and they activate liver X receptors (LXRs), which promote cholesterol efflux, modulate inflammatory responses, and protect mice during Mtb infection.^27–30^ Oxysterols thus link macrophage lipid homeostasis to host defense, yet whether Mtb directly manipulates this oxysterol-LXR axis is unknown.

We previously identified cholest-4-en-3-one, a metabolite generated from cholesterol, as elevated in sputum from patients with active TB.^31^ Cholest-4-en-3-one is produced by the Mtb enzyme 3β-hydroxysteroid dehydrogenase (3β-Hsd), which oxidizes the 3β-hydroxyl group of sterols and isomerizes the Δ^5^ double bond to Δ^4^, generating 3-oxo-Δ^4^ metabolites (Fig. 1A).^32^ Unlike mammalian 3β-HSD enzymes, which generally act on a narrow range of steroid substrates,^33,34^ recombinant Mtb 3β-Hsd metabolizes a broad range of sterols,^32,35^ raising the question of whether Mtb modifies oxysterols with immunometabolic functions. To profile sputum oxysterols from geographically distinct cohorts, we used targeted liquid chromatography–mass spectrometry with multistage fragmentation (LC–MS(MS^3^)). We identified a TB-associated oxysterol signature characterized by the conversion of 27-HC and 3β-HCA to 3-oxo-Δ^4^ metabolites [(27-hydroxycholest-4-en-3-one (27-HCO) and 3-oxocholest-4-enoic acid (3O-CA)]. We further showed, in an independent geographical cohort using a distinct liquid chromatography tandem mass spectrometry (LC-MS/MS) platform, that 27-HCO and 3O-CA declined in TB patients after two months of treatment. Unlike the parent compounds, 27-HCO and 3O-CA failed to activate LXR target-gene expression, and 3O-CA lost its ability to inhibit the SREBP2 pathway. Within infected macrophages, the conversion of 27-HC to 27-HCO required Mtb 3β-Hsd, which promoted macrophage cholesterol accumulation. Together, these findings identify CYP27A1*-*3β-Hsd co-metabolites as candidate TB biomarkers and reveal a previously unrecognized bacterial strategy: Mtb enzymatically inactivates endogenous oxysterols to disrupt macrophage cholesterol homeostasis, thereby generating disease-associated oxysterol signatures.

**Fig. 1.**
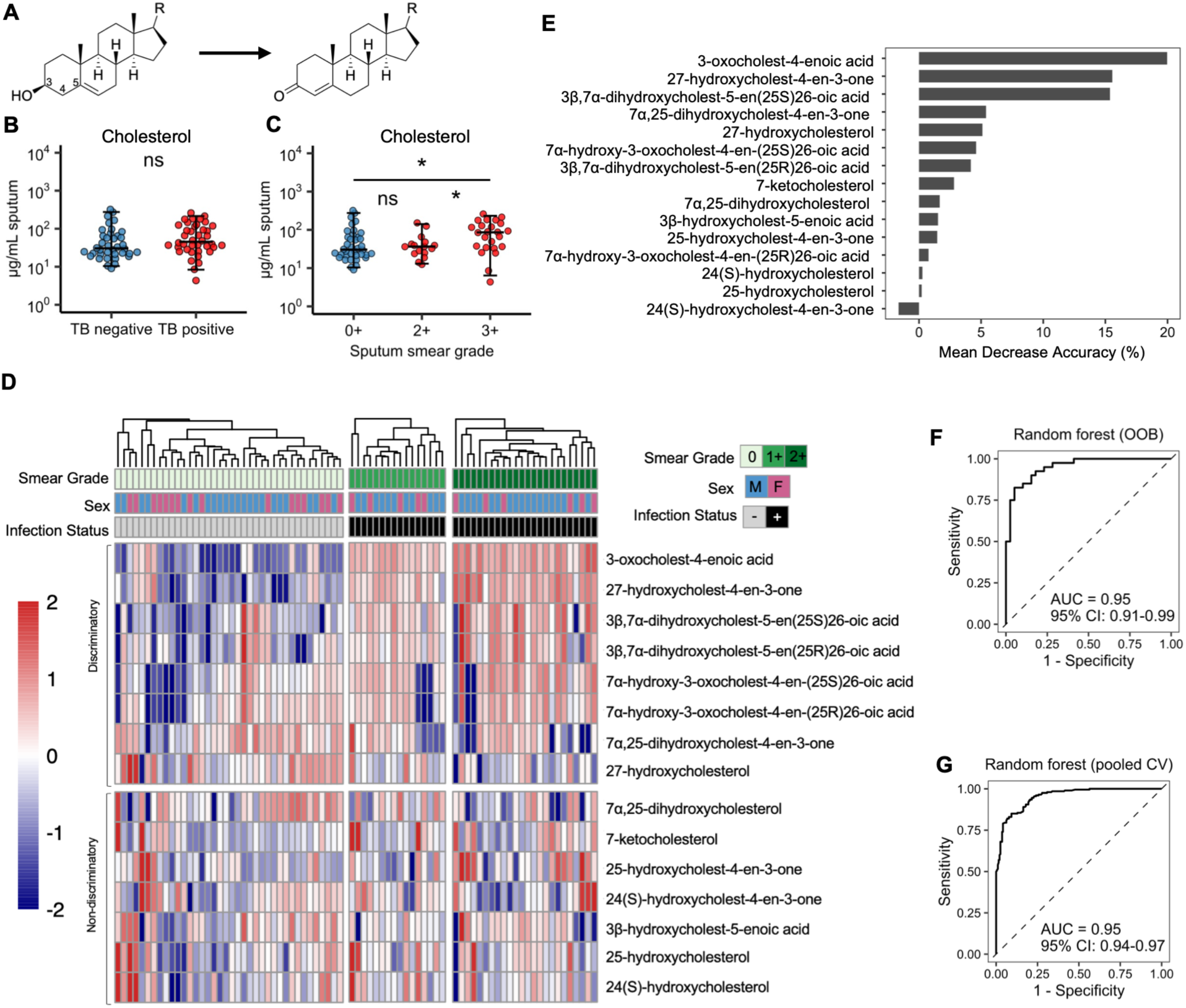
TB creates a distinct oxysterol landscape in the lungs. (**A**) The Mtb enzyme 3β-Hsd oxidizes the 3β-hydroxyl group of sterols and isomerizes the Δ^5^ bond. (**B-D**) Sputum oxysterol concentrations from 78 subjects were quantified using LC-MS(MS^3^). Sputum free (non-esterified) cholesterol levels were stratified by (**B**) infection status and (**C**) smear grade. (**D**) Sputum oxysterol concentrations from 78 subjects underwent normalization to free cholesterol, log10 transformation, and z-scoring. Samples are stratified by sputum smear grade horizontally and by ability to discrimination of TB status on ROC analysis vertically. Top panel shows oxysterols that were individually predictive of TB status using ROC analysis. Columns were clustered based upon the discriminative oxysterols. (**E**) Variable importance plot from random forest classifier trained using the oxysterols in (**D**). (**F-G**) ROC curves from out-of-bag (OOB) and pooled cross validation (CV) predictions of 78 subjects, with model performance indicated by AUC and 95% confidence interval (CI). Statistical significance was assessed using Wilcoxon rank sum tests (**B**) and Kruskal– Wallis tests with Dunn’s post hoc correction for multiple comparisons (**C**). Bars represent median, and whiskers encompass 95% of the observed data. * P < 0.05; NS, not significant. Data from individual metabolites are also shown in Fig. 2, Fig. S2, and Fig. S3.

## Results

### Active TB creates a distinctive oxysterol landscape in the lungs

To define whether TB reshapes the pulmonary oxysterol repertoire, we analyzed sputum from 78 HIV-negative adults with TB or TB-like symptoms in Peru and Vietnam (Table S1, Table S2).^31^ Forty individuals had microbiologically confirmed active TB, and 38 symptomatic control individuals had negative sputum smear microscopy, culture, GeneXpert, and QuantiFERON-TB Gold, indicating an alternative diagnosis. Age and sex distribution did not differ significantly between TB-positive and TB-negative individuals (Table S2). After sputum processing, we quantified free (non-esterified) cholesterol and 15 oxysterol species using LC–MS(MS^3^) (Fig. S1). Cholesterol concentrations trended higher in TB-positive versus TB-negative individuals, with a statistically significant increase in patients with the highest mycobacterial burden (smear grade 3+) compared with TB-negative controls (Fig. 1B–C). After normalization to free cholesterol levels in the sputum, eight oxysterols significantly discriminated TB-positive from TB-negative sputum (Fig. 1D). Among these metabolites, only 27-HC and 7α,25-dihydroxycholest-4-en-3-one (7α,25-diHCO) were decreased in the TB-positive samples, whereas six metabolites were increased. A random forest classifier based on sputum oxysterol concentrations distinguished TB-positive from TB-negative participants and identified the 3-oxo-Δ^4^ oxysterols 27-HCO and 3O-CA as the strongest model predictors (Fig. 1E). The model had an out-of-bag area under the receiver operating characteristic curve (AUROC) of 0.95 (95% confidence interval (CI), 0.91-0.99); cross-validated predictions also yielded an AUROC of 0.95 (95% CI, 0.94-0.97) (Fig. 1F-G). Thus, active TB is associated with a distinctive sputum oxysterol signature characterized by reduced 27-HC and increased 3-oxo-Δ^4^ metabolites.

### 27-HCO and 3O-CA are enriched in sputum during active TB

Seven of the eight discriminative oxysterols identified in the univariate analysis are metabolites derived from CYP27A1 products. CYP27A1 generates 27-HC and subsequently 3β-HCA (Fig. 2A), which gives rise to additional oxidized species that are intermediates in bile acid synthesis (Fig. S2A). 27-HCO and 3O-CA are 3-oxo-Δ^4^-derivatives of 27-HC and 3β-HCA, respectively (Fig. 2A). Given that 27-HCO and 3O-CA had the highest discriminatory power, and they are not well-described human metabolites, we analyzed them and their immediate precursors in more detail (Fig. 2B-U). Actual and cholesterol-normalized sputum concentrations of 27-HCO and 3O-CA were increased in participants with TB and were highest among those with smear grade 3+ disease, consistent with an association with mycobacterial burden (Fig. 2B-M). Most TB-negative individuals had sputum concentrations of 3O-CA below the limit of detection (0.01 ng/mL), while the median concentration in TB-positive individuals was 3.76 ng/mL (Fig. 2E). While actual sputum 27-HC concentrations were similar between groups (Fig. 2B), 27-HC was significantly lower in TB-positive sputum following normalization to cholesterol (Fig. 2F), with the largest reductions seen in subjects with the highest smear grade (Fig. 2J). Both 27-HCO and 3O-CA were highly discriminative between groups, with AUROC of 0.92 and 0.95, respectively (Fig. 2O, 2Q). 27-HC yielded an AUROC of 0.32 (95% CI 0.20-0.44), reflecting an inverse association with active TB (Fig. 2N). From these same subjects, we next measured free cholesterol and oxysterols in the plasma (Fig. 2R-U; Fig. S2B). We found that plasma 27-HC was significantly reduced in TB-positive plasma, mirroring our finding in sputum (Fig. 2R). There were no significant differences in the median concentrations of 27-HCO and 3O-CA in plasma between TB and control individuals (Fig. 2S, 2U). These findings identify a pulmonary TB-associated oxysterol signature characterized by depletion of 27-HC and accumulation of 27-HCO and 3O-CA, a pattern consistent with conversion of endogenous CYP27A1-derived oxysterols to 3-oxo-Δ^4^ oxysterol metabolites in infected lung.

**Fig. 2.**
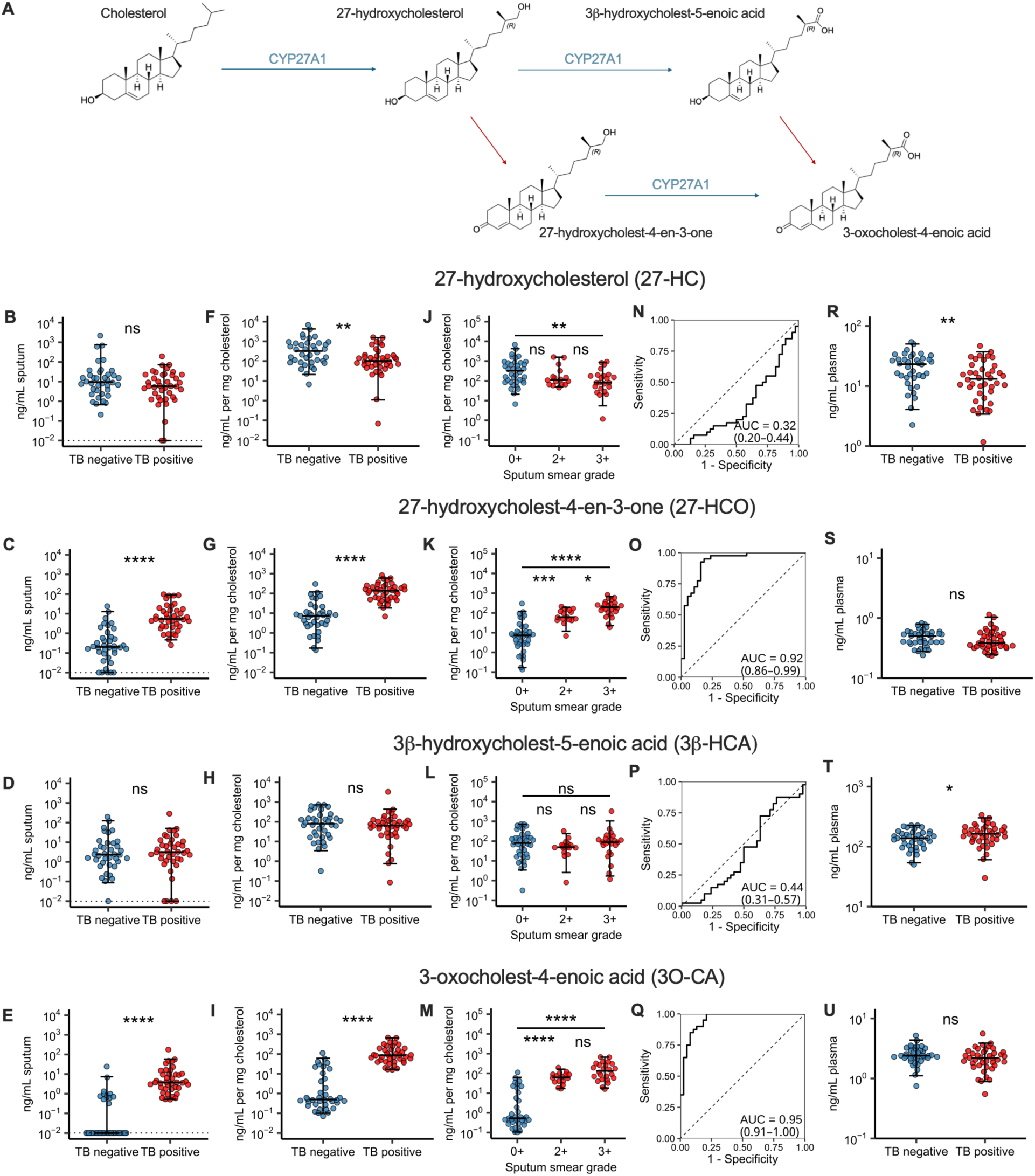
27-hydroxycholest-4-en-3-one and 3-oxocholest-4-enoic acid are elevated in sputum during TB and predict TB status. (**A**) Model of host-bacterial co-metabolism during TB. Host CYP27A1 sequentially oxidizes cholesterol to 27-HC and 3β-HCA. Sputum and plasma oxysterol levels from 78 subjects were quantified using LC-MS(MS^3^). Sputum oxysterol concentrations (**B-E**), sputum oxysterol concentrations normalized to free cholesterol (**F-I**), normalized oxysterol concentrations stratified by smear grade (J-M), ROC curves (**N-Q**), showing ability of each normalized oxysterol concentration to predict TB status, and plasma oxysterol concentrations (**R-U**). Statistical significance was assessed using Wilcoxon rank-sum tests (**B-I**; **R-U**) and Kruskal– Wallis tests with Dunn’s post hoc correction for multiple comparisons (**J-M**). Bars represent median and whiskers encompass 95% of the observed data. (**B-E**) The limit of detection is indicated by the dotted line (**B-E**). 95% confidence intervals of ROC curves estimated by the DeLong method (**N-Q**). **** P < 0.0001; *** P < 0.001; ** P < 0.01; * P < 0.05; NS, not significant.

Four additional oxysterols that discriminated TB-positive from TB-negative sputum were 7α-substituted cholestenoic acids that are generated from 3β-HCA (Fig. S2), including the 25R and 25S epimers of 3β,7α-dihydroxycholest-5-enoic acid (3β,7α-diHCA) and 7α-hydroxy-3-oxocholest-4-enoic acid (7αH,3O-CA) (Fig. S2C-M). The 7α-substituted cholestenoic acid metabolites showed moderate discriminatory performance for active TB (AUC 0.63-0.85). Plasma concentrations of 3β-HCA and both 3β,7α-diHCA epimers were also increased in individuals with TB (Fig. 2T, Fig. S2N, P). These findings indicate a broad perturbation of CYP27A1-derived cholestenoic acid metabolism during TB, with the most marked differences seen in 27-HCO and 3O-CA levels in the lungs.

In contrast to the CYP27A1-derived metabolites, the median sputum concentrations of 25-HC, 25-hydroxycholest-4-en-3-one (25-HCO), 7α,25-dihydroxycholesterol (7α,25-diHC), 24(S)-hydroxycholesterol [24(S)-HC], 24(S)-hydroxycholest-4-en-3-one [24(S)-HCO], and 7-ketocholesterol (a common autoxidation product) did not differ significantly between TB-positive and TB-negative individuals following normalization to free cholesterol (Fig. S3). After normalization to cholesterol, sputum 7α,25-diHCO was lower in TB-positive compared to TB-negative sputum (Fig. S3R). Plasma 25-HC, 7α,25-diHC, 7α,25-diHCO, and 7-ketocholesterol also did not differ between groups (Fig. S3AD-AG). In subjects with TB, plasma 24(S)-HC, which is derived primarily from the central nervous system (CNS), was lower. Importantly, we noted only minor differences in sputum oxysterol concentrations between participants from Peru or Vietnam (Table S1). Thus, the active TB oxysterol signature was dominated by changes in CYP27A1-derived metabolites in sputum, with only limited additional alterations in sputum 7α,25-diHCO and circulating CNS-derived 24(S)-HC, rather than a broad perturbation of sterol oxidation.

### Cholest-4-en-3-one, 27-HCO, and 3O-CA decline in sputum with TB treatment

Having demonstrated enrichment of 27-HCO and 3O-CA in TB sputum relative to symptomatic controls across two geographically diverse cohorts, we next asked how these oxysterols respond to TB therapy. To provide orthogonal analytical validation, we quantified the oxysterols using a distinct LC-MS/MS platform that did not involve a derivatization step. We analyzed paired sputum samples collected before treatment and after 2 months of standard therapy from 30 HIV-negative adults with drug-susceptible pulmonary TB in Uganda and Kenya (Fig. 3A-M, Table S1); all participants were cured after 6 months of therapy. This cohort included 15 male and 15 female participants selected at random from successfully treated individuals; 80% had cavitary disease. After 2 months of treatment, total, free, and esterified cholesterol in the sputum declined (Fig. 3A-C). Actual and cholesterol-normalized concentrations of 27-HCO and 3O-CA also markedly decreased after treatment (Fig. 3F, 3H, 3K, 3M). In all treated participants, 27-HCO declined from a median of 27.2 ng/mL before treatment to below the lower limit of detection (0.1 ng/mL) after treatment (Fig. 3F). Similarly, the median concentration of 3O-CA before treatment was 17.3 ng/mL and became undetectable in 66.6% of participants after 2 months of treatment (Fig. 3H). In contrast, 27-HC and 3β-HCA did not differ significantly before and after treatment (Fig. 3E, 3G). We also measured cholest-4-en-3-one, an Mtb-derived cholesterol metabolite that we previously showed is associated with TB status.^31^ Although cholest-4-en-3-one significantly declined after treatment, substantial levels persisted after 2 months in contrast to 27-HCO and 3O-CA (Fig. 3D, 3I). Thus, in a geographically independent cohort analyzed by an alternative LC-MS/MS workflow, sputum 27-HCO and 3O-CA declined, often to undetectable levels, supporting their potential use as biomarkers of active pulmonary TB and treatment response.

**Fig. 3.**
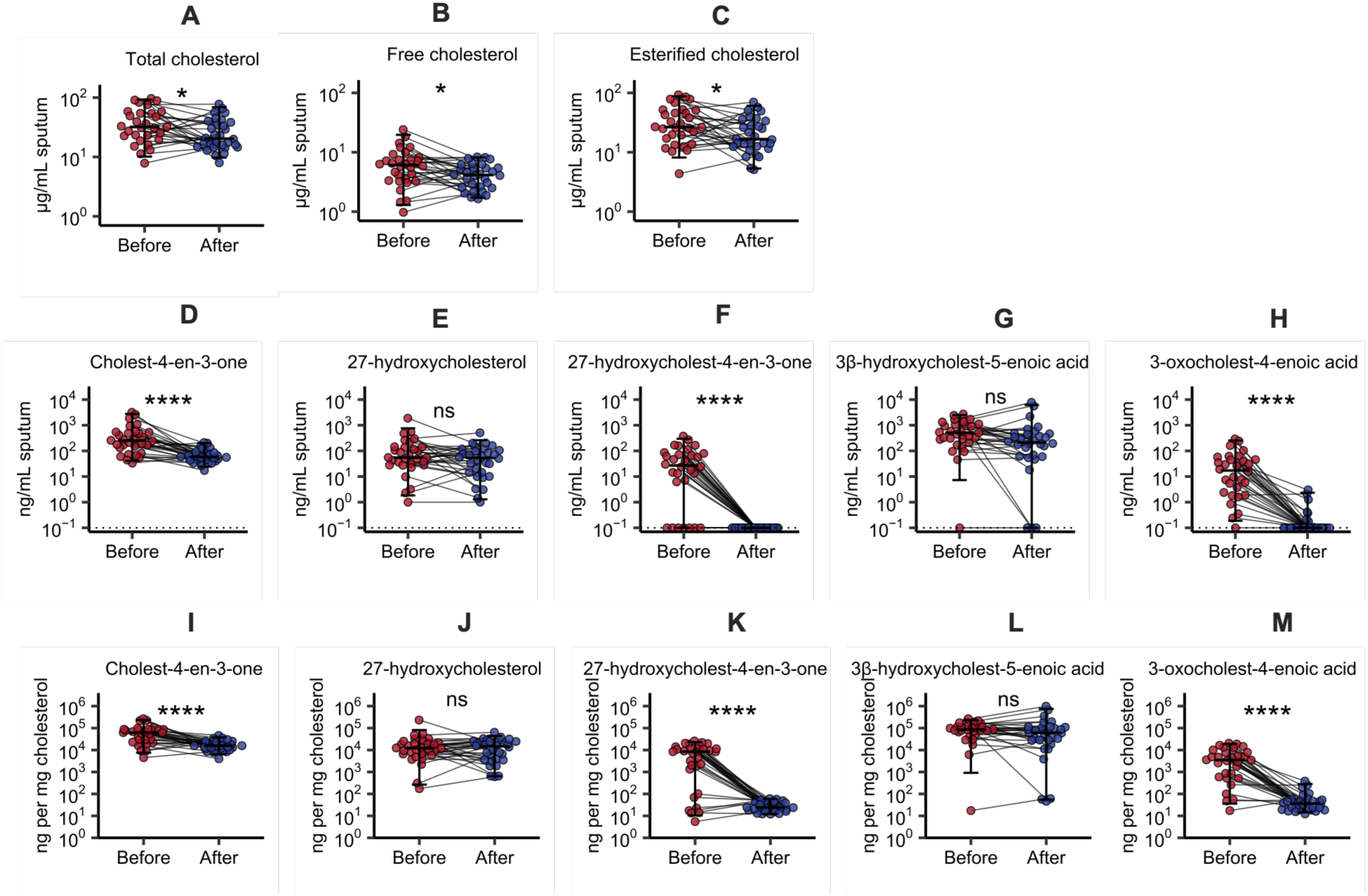
Sputum concentrations of cholest-4-en-3-one, 27-hydroxycholest-4-en-3-one, and 3-oxocholest-4-enoic acid decline with TB treatment. **(A-M)** Cholesterol and oxysterol concentrations were measured using LC-MS/MS from paired sputum samples from participants before initiation of anti-tuberculous therapy (“Before”) and after 8 weeks of antibiotic treatment (“After”). **(D-H)** Actual oxysterol concentrations and **(I-M)** normalized to free cholesterol. The limit of detection is indicated by the dotted line **(D-H)**. Bars represent median and whiskers encompass 95% of the observed data. Statistical significance was assessed using the Wilcoxon matched pairs signed rank test; **** P < 0.0001; * P < 0.05; NS, not significant.

### Mtb 3β-Hsd generates 3-oxo-Δ^4^ metabolites during infection

To identify cellular sources of the CYP27A1-derived oxysterols in TB, we reanalyzed a publicly available single-cell RNA sequencing (scRNAseq) dataset generated from lung tissue of individuals recently treated for TB and uninvolved lung tissue from TB-negative individuals who underwent resection for lung malignancy.^36^ Monocyte-derived macrophages (MDM) and alveolar macrophages (AM) were the predominant cell types expressing *CYP27A1* (Fig. 4A, Fig. S4A). Although per-cell *CYP27A1* expression was comparable between groups, lungs from individuals with recent TB contained more CYP27A1-expressing macrophages, resulting in higher overall *CYP27A1* expression. We also examined *CH25H* and *CYP46A1*, which encode enzymes that generate 25-HC and 24(S)-HC, respectively. *CYP46A1* expression was undetectable, whereas *CH25H* expression was modest and comparable between groups, consistent with the lack of TB-associated difference in sputum 25-HC, 24(S)-HC, and 7α,25-diHC levels (Fig. S4B). These findings identify macrophages, a major cellular niche of Mtb, as the principal lung cell population expressing CYP27A1 and hence the likely sources of 27-HC and 3β-HCA. The mechanism generating the corresponding 3-oxo-Δ^4^ metabolites, 27-HCO and 3O-CA, is less clear because no candidate human enzyme is expressed in the relevant lung cell populations. HSD3B7 has a well-defined role metabolizing 7α-substituted cholestenoic acids, but HSD3B7 requires that its substrates have a 7α-hydroxy group, which is lacking in 27-HC and 3β-HCA. HSD3B1 and HSD3B2 primarily oxidize steroids that lack the extended side chain that is present in 27-HC and 3β-HCA. Although HSD3B1 can oxidize oxysterols as minor products,^34^ HSD3B1 and HSD3B2 are expressed predominantly in placental tissue, skin, and adrenal-gonadal tissue. In a scRNAseq dataset of 107 healthy human lungs, only 0.0017% of cells express HSD3B1 and 0.0032% of cells express HSD3B2, with the primary expressors being mesothelial cells and fibroblasts (Human Lung Cell Atlas; CZ CELLxGENE Discover).^37^ HSD3B1 and HSD3B2 were not detected in major lung cell populations in individuals with recent TB in the Mbano *et al.* dataset. These findings argue against a host 3β-HSD as the source of the 27-HCO and 3O-CA detected in TB sputum. Because recombinant Mtb 3β-Hsd can oxidize sterols irrespective of 7α-substitution,^32,35^ we hypothesized that 3β-Hsd generates 27-HCO and 3O-CA during infection. To test this, we infected MDMs from healthy donors with Mtb lacking 3β-Hsd (Mtb Δ*hsd*) or a complemented strain expressing *hsd* from its native promoter (Mtb Δ*hsd::hsd*). Cholest-4-en-3-one and 27-HCO were significantly higher in the cultures infected with 3β-Hsd-expressing Mtb compared to uninfected cells and those infected with Δ*hsd* Mtb (Fig. 4BC). In most infected and uninfected samples, 3O-CA was below the limit of detection (10 pg/10^6^ cells), which may be because 3β-HCA was found at ∼ 20-fold lower concentrations than 27-HC (Fig. S4CD). While 25-HC was present (Fig. S4CE), its 3-oxo-Δ^4^ metabolite (25-HCO) was below the limit of detection (10 pg/10^6^ cells). Thus, in macrophages, Mtb 3β-Hsd promotes the generation of the 3-oxo-Δ^4^ metabolite of 27-HC.

**Fig. 4.**
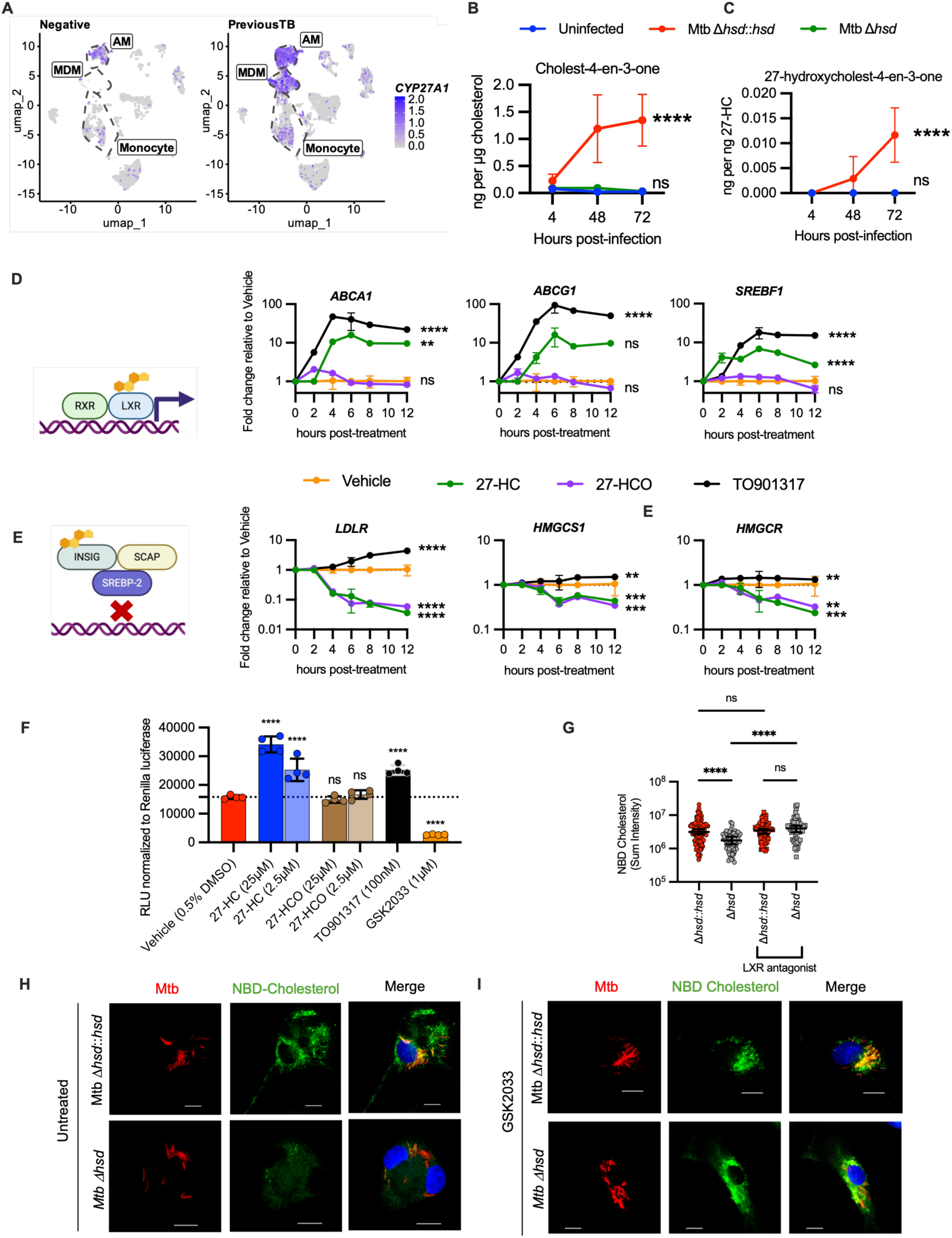
Mtb 3β-Hsd promotes 27-HCO formation and impairs macrophage cholesterol homeostasis. (**A**) Expression of *CYP27A1* from scRNA-seq of lung resections from TB-negative individuals and those with recent TB. Alveolar macrophage (AM), monocyte-derived macrophage (MDM), and monocytes are designated by dotted lines. (**B-C**) Human MDMs from 4 human donors were infected with Δ*hsd* Mtb, Δ*hsd*::*hsd* Mtb, or uninfected. Cholesterol concentration was calculated using Amplex Red Cholesterol Assay Kit and oxysterol concentrations were quantified by LC-MS/MS. Data are expressed as a ratio of 3-oxo-Δ^4^ metabolite to the corresponding 3β-hydroxy metabolite at indicated time points post-infection. (**D-E**) THP-1 macrophages were treated with 27-HC (5 µM), 27-HCO (5 µM), T0901317 (100 nM), or vehicle (0.25% DMSO), and cells were harvested 2, 4, 6, 8, and 12 h after treatment. Expression is shown as fold change relative to vehicle-treated controls. Data are representative of 3 independent experiments. (**F**) HEK293 cells expressing the LXR response element–firefly luciferase reporter, LXRα, and Renilla luciferase (transfection control) were treated for 24 hours with vehicle, 27-HC, 27-HCO, the LXR agonist T0901317, or LXR antagonist GSK2033 at indicated dose. Firefly luciferase relative light units (RLU) were normalized to Renilla luciferase activity for each well. (**G-I**) Human MDMs were infected with Δ*hsd* or Δ*hsd*::*hsd* at MOI of 5 and treated with vehicle or LXR antagonist GSK2033 (1µM) for 48 h. NBD-cholesterol was added 3 h prior to harvest. (**G**) Sum intensity of NBD-cholesterol from individual cells. (**H-I**) Representative immunofluorescence images. Two-way ANOVA with Dunnett’s multiple comparisons test of each treatment versus vehicle control (**B-E**), one-way ANOVA with Dunnett’s multiple comparisons test of each treatment versus vehicle control (**F**), and Kruskal-Wallis test with Dunn’s correction (**G**). Error bars in (**B-G**) show mean and standard deviation. **** P < 0.0001; *** P < 0.001; ** P < 0.01; NS, not significant.

### 27-HCO and 3O-CA fail to activate LXR-dependent responses and 3O-CA fails to suppress SREBP2 target gene expression

27-HC is a major lung oxysterol that contributes to cholesterol homeostasis through separable mechanisms: it can activate LXR to promote cholesterol efflux, and it can suppress the SREBP2 pathway through INSIG-dependent ER retention of the SCAP-SREBP2 complex, thereby reducing cholesterol uptake and synthesis (Fig. 4DE).^24,38,39^ Since Mtb converts 27-HC to 27-HCO, we considered that this reaction might alter the signaling properties of 27-HC and thereby rewire host cholesterol homeostasis. To test this possibility, we treated THP-1 macrophages with 27-HC, 27-HCO, the synthetic LXR agonist TO901317, or vehicle and quantified LXR- and SREBP2-responsive transcripts. Treatment with 27-HC, but not 27-HCO, induced transcription of the LXR target genes ATP binding cassette transporter A1 (*ABCA1*), ATP binding cassette transporter G1 (*ABCG1*), and sterol regulatory element binding transcription factor 1 (*SREBF1*) (Fig. 4D). In contrast, both 27-HC and 27-HCO similarly repressed the SREBP2 target genes low-density lipoprotein receptor (*LDLR*), HMG-CoA synthase 1 (*HMGCS1*), and HMG-CoA reductase (*HMGCR*) (Fig. 4E), indicating that oxidation of 27-HC selectively abolishes its LXR agonist activity while preserving suppression of SREBP2. We next used a luciferase reporter assay in which LXRα activation of an LXR-response element drives firefly luciferase expression. 27-HC activated the reporter in a dose-dependent manner, whereas 27-HCO did not (Fig. 4F). Thus, conversion of 27-HC to 27-HCO eliminates LXR agonism, but both compounds inhibit the SREBP2 pathway. We next examined 3β-HCA, which is a comparatively weak LXR agonist,^30^ and its 3-oxo-Δ^4^ metabolite, 3O-CA. 3β-HCA modestly and transiently induced expression of the LXR target genes *ABCA1*, *ABCG1*, and *SREBF1* (Fig. S4F). In contrast, 3O-CA did not induce expression of the canonical LXR-responsive genes *ABCA1* or *ABCG1,* indicating, as we found with 27-HC, that conversion to the 3-oxo-Δ^4^ metabolite disrupted the ability of 3β-HCA to activate LXR-dependent gene expression. However, in contrast to both 27-HC and 27-HCO being able to suppress SREBP2 target genes, 3O-CA failed to inhibit *LDLR*, *HMGCS1,* and *HMGCR* as we saw for the parent compound, 3β-HCA (Fig. S4G). Together, these findings show that in humans with TB the major lung LXR agonist, 27-HC, is converted into a metabolite with diminished LXR activity. In addition, 3β-HCA is converted to a metabolite with diminished ability to inhibit SREBP2-dependent gene expression.

### Mtb 3β-Hsd alters macrophage cholesterol handling

Since Mtb inactivated the LXR and SREBP2 signaling properties of 27-HC and 3β-HCA, we considered whether 3β-Hsd alters macrophage cholesterol homeostasis. To test this, we infected human MDMs with mCherry-expressing Mtb Δ*hsd* or the complemented strain (Mtb Δ*hsd::hsd*). After 48 hours of infection, MDMs were incubated with 22-nitrobenzoxadiazole-cholesterol (NBD-cholesterol), a fluorescent cholesterol analogue. Macrophages infected with the 3β-Hsd-complemented strain exhibited significantly more intracellular retention of NBD-cholesterol than macrophages infected with Mtb Δ*hsd* (Fig. 4GH). To determine if the lower levels of NBD-cholesterol seen during infection with Mtb Δ*hsd* was LXR-dependent, MDMs were cultured in medium containing the LXR antagonist GSK2033 or vehicle control during the infection. Pharmacologic LXR inhibition restored NBD-cholesterol retention during Mtb Δ*hsd* infection, indicating that 3β-Hsd impairs LXR-dependent remodeling of macrophage cholesterol handling (Fig. 4G-I).

## Discussion

Mtb can use host derived cholesterol as a source of carbon and energy, and cholesterol utilization contributes to bacterial persistence during chronic infection.^15^ Recent work demonstrates that macrophage cholesterol accumulation can impair antigen presentation and protective CD4+ T cell responses.^9-13,15,16,40^ Mycobacterial cell envelope lipids, including oxygenated mycolic acids, have been implicated in remodeling macrophage lipid metabolism, and in some contexts, promoting cholesteryl ester accumulation;^13,41^ however, the mechanisms by which Mtb perturbs host cholesterol homeostasis remain incompletely defined. Here, we identify a host-pathogen co-metabolic pathway through which Mtb reshapes macrophage cholesterol metabolism. Across three cohorts from three continents, sputum from individuals with TB was characterized by accumulation of the 3-oxo-Δ^4^ sterols, 27-HCO and 3O-CA, together with a reduction in 27-HC relative to cholesterol. The CYP27A1-derived oxysterol axis may be particularly susceptible to modification by Mtb because CYP27A1 is highly expressed in lung macrophages.^36^ 27-HCO is normally only a minor human metabolite,^34,42–44^ and we found that it was generated in Mtb-infected macrophages in a 3β-Hsd-dependent manner. Conversion of 27-HC and 3β-HCA to their 3-oxo- Δ^4^ metabolites abrogated LXR signaling by both molecules, while also disrupting suppression of SREBP2 by 3β-HCA. These findings establish a mechanism by which Mtb enzymatically disrupts pathways governing macrophage cholesterol homeostasis.

These findings reveal an unusual form of host-pathogen metabolic interaction. Rather than solely consuming cholesterol as a nutrient or indirectly altering host lipid pathways, Mtb enzymatically modifies endogenous signaling molecules that govern cholesterol homeostasis. Oxidation of 27-HC abolished its LXR agonist activity while preserving its capacity to suppress SREBP2-responsive genes, whereas oxidation of 3β-HCA disrupted both activities. Consistent with these signaling effects, macrophages infected with 3β-HSD-expressing Mtb retained more NBD-cholesterol than those infected with Δ *hsd* Mtb, and pharmacologic LXR inhibition eliminated this difference. Thus, a single bacterial enzyme uncouples complementary arms of host sterol regulation, with the net effect of increasing cholesterol retention in infected macrophages. More broadly, pathogen-mediated chemical modification of host metabolic signals may represent an underrecognized strategy for remodeling the intracellular environment.

Our findings establish that 3β-Hsd-dependent oxysterol oxidation alters LXR-dependent cholesterol handling in human macrophages, but they do not establish that this metabolic rewiring confers a direct bacterial fitness advantage. Indeed, our previous work showed that deletion of *hsd* did not produce a detectable growth defect in macrophages under the conditions examined.^31^ The consequences of oxysterol inactivation may therefore be context-dependent, reflecting effects of cholesterol availability, macrophage function, or immune interactions that are not captured by bacterial growth assays in isolated macrophages. Defining how this pathway influences persistence, immune control, and granuloma biology will require experimental systems that reproduce the cellular complexity and oxysterol environment of the human lung.

The metabolites we identified have potential clinical utility. Cholest-4-en-3-one, 27-HCO, and 3O-CA declined after eight weeks of successful TB treatment, supporting their potential use as treatment-responsive biomarkers. 27-HCO and 3O-CA showed greater discrimination between TB cases and controls than cholest-4-en-3-one, which has appreciable baseline abundance in TB-negative individuals and in individuals after two months of treatment, consistent with additional host or microbial sources. The near absence of 3O-CA in most TB-negative samples suggests that it may be especially informative. However, its specificity should be evaluated in populations in which alternative sources of 3-oxo-Δ^4^ metabolites may be present, including pregnant individuals, in whom placental HSD3B1 is highly expressed^34^ and in patients with nontuberculous mycobacterial (NTM) lung disease, because 3β-Hsd is conserved in NTMs. The reduction of 27-HCO and 3O-CA during therapy is consistent with their dependence upon active bacterial metabolism, but the present study was not designed to define their relationship to time to culture conversion, residual bacterial burden, relapse, or treatment failure. Prospective studies incorporating serial sputum microbiology in clinically diverse populations will be needed to determine the clinical utility of these potential biomarkers. Because a non-sputum-based diagnostic is a major clinical need, the modest discriminatory performance of plasma cholestenoic acids should be evaluated in combination with other circulating biomarkers. The diagnostic performance of the metabolites we identified should be evaluated in individuals with low-burden disease and in additional clinically important populations, including people living with HIV and those with diabetes.

The oxysterol pattern observed in pulmonary TB differs from that described in atherosclerosis. In atherosclerotic lesions, 27-HC accumulates markedly relative to non-diseased vasculature, consistent with a compensatory response to cellular cholesterol loading.^45,46^ By contrast, 27-HC was reduced in TB, while its oxidized product accumulated. This pattern supports a model in which Mtb actively diverts 27-HC to promote foam cell formation, rather than low 27-HC levels simply reflecting a generic consequence of foam cell formation. In addition, we identified elevated levels of several cholestenoic acid and bile acid pathway intermediates in sputum and plasma from individuals with TB. These changes might reflect compensatory mechanisms to dispose of cholesterol, increased numbers of macrophages in the lungs, or additional host–pathogen metabolic interactions. The biological consequences of these changes require further study. In addition, recombinant Mtb 3β-Hsd has been shown to metabolize a broad array of substrates, including androgen precursors,^32,35^ suggesting its impact may extend beyond CYP27A1-derived oxysterol metabolism.

LXR is an important component of host defense against Mtb. In addition to promoting cholesterol efflux, LXR also regulates the inflammatory response, and its activation is protective in macrophage and mouse models of Mtb infection.^27,28,47^ Consistent with this role, genetic variation in NR1H3 and NR1H2, which encode LXRα and LXRβ, respectively, has been associated with TB susceptibility.^48^ We show that Mtb reduces the level of a major pulmonary LXR agonist 27-HC and increases 27-HCO, reducing the 27-HC:27-HCO ratio by approximately 100-fold in individuals with active TB. These data identify LXR signaling as a point of host-pathogen conflict and suggest that restoring LXR activity could represent a host-directed therapeutic strategy. Because treatment with systemic administration of LXR agonists can promote hepatic steatosis,^49^ targeted pulmonary delivery or approaches that prevent local oxysterol inactivation may offer a tractable route for therapeutic development.

## Resource availability

### Lead contacts

Further information and requests for resources and reagents should be directed to and will be fulfilled by the lead contact Jennifer Philips.

### Materials availability

all materials generated in this study will be made available from the lead contact upon request.

### Data and code availability

all data used in the analysis will be made available. This publication does not report original code.

## Acknowledgements

Research reported in this publication was supported by the Washington University Institute of Clinical and Translational Sciences grant UL1TR002345 from the National Center for Advancing Translational Sciences of the National Institutes of Health (NIH). The content is solely the responsibility of the authors and does not necessarily represent the official view of the NIH or other funders. This work was made possible by FIND and TB Alliance who shared clinical samples. We thank Morten Ruhwald and Kavi Ramjeet at FIND and Majda Benhayoun at TB Alliance for their assistance. Funding was provided by the National Institutes of Health grant AI178685-01A1 (JAP); National Institutes of Health grant KL2TR002346 (ATR); Washington University School of Medicine (JAP); Biotechnology and Biological Sciences Research Council - BBSRC (BB/L001942/1 – YW, WJG, BB/S019588/1 – WJG, YW); Medical Research Council MRC (MR/X012387/1 – WJG, YW); European Union through European Structural Funds (ESF), as part of the Welsh Government funded Academic Expertise for Business project (WJG, YW).

## Author contributions

Conceptualization: JAP, ATR, WJG, YW

Methodology: XJ, WJG, YW, DFC

Investigation: ATR, MAA, PC, JM, AL, SB, IAW, MQ

Formal Analysis: ATR, MAA, PC, JM, AL, IAW, CWG, DK, WJG, YW, JAP

Visualization: ATR, PC, DK

Funding acquisition: JAP, WJG, YW

Project administration: JAP, WJG, YW

Supervision: JAP, WJG, YW, DC, BTE Writing – original draft: ATR

Writing – review & editing: ATR, MAA, PC, SB, BTE, DK, DFC, WJG, YW, JAP

## Star Methods

### Bacterial strains

*Mycobacterium tuberculosis* (H37Rv strain) was grown under aerobic conditions at 37°C in Middlebrook 7H9 broth with 0.05% Tyloxapol, 0.2% glycerol, and 10% OADC (oleic acid, albumin, dextrose, catalase; BD Biosciences). Δ*hsd* Mtb and the complemented strain (Δ*hsd::hsd* pCH89) were previously reported.^31^ For immunofluorescence experiments, strains constitutively expressed mCherry from an episomal plasmid under the control of the smyc promoter. Hygromycin (50µg/mL) and Zeocin (Invivogen, catalog no. ant-zn-05; 25µg/mL) were added when required for plasmid selection.

### Cell culture

Human monocyte-derived macrophages (MDM) were differentiated from peripheral blood mononuclear cells obtained from healthy donors using RPMI supplemented with 10% FBS, 2 mM L-glutamine, and 50 ng/mL human M-CSF (Gibco, Thermo Fisher Scientific, catalog no. 300-25) for 7 to 10 days. Following harvest, MDMs were allowed to adhere overnight prior to infection or treatment. HEK293TN cells were purchased from ATCC and cultured in DMEM supplemented with 10% FBS, non-essential amino acids, and 25 mM HEPES. Cells were allowed to adhere overnight prior to treatments. THP-1 cells were obtained from ATCC and cultured in RPMI supplemented with 10% FBS, 2 mM L-glutamine, and 55 µM β-mercaptoethanol. THP-1 cells were differentiated by incubating with 20ng/mL phorbol 12-myristate 13-acetate (PMA) for 24 hours. On the following day, cells were washed to remove PMA and rested for 24 hours before use in experiments.

### Clinical samples and study approval

Samples from people with and without TB from Peru and Vietnam were obtained from the Foundation for Innovative New Diagnostics (FIND; https://www.finddx.org/specimen-bank/specimens-tb/). 78 of the previously described 80 samples were analyzed ^31^. Two samples were unable to be analyzed due to inadequate remaining sputum. Samples from individuals on TB treatment were provided by the Measurement and Assay Resource for Tuberculosis (MARK-TB), a resource founded by the Tuberculosis Trials Consortium (TBTC). Samples were provided from TBTC Study 36 (“Platform for Assessment of TB Treatment Outcomes”), an observational, multicenter study sponsored by the U.S. CDC, in which participants received standard of care antibiotic therapy for drug-susceptible pulmonary TB. All specimens were collected at qualified clinics under a protocol approved by an Institutional Review Board and obtained with informed consent. FIND and MARK-TB follow good clinical and laboratory practice in obtaining and processing sample, using standardized procedures that maximize the reliability of diagnostic evaluations.

### Chemicals and reagents

The following compounds were used as internal standards or reference compounds: [2,2,4,6,6-^2^H_5_]cholest-4-en-3-one (MedChemExpress, catalog no. HY-113365S), [25,26,26,27,27,27-^2^H_6_](25R)26-hydroxycholesterol (Avanti Polar Lipids, catalog no. A80059), [25,26,26,27,27,27-^2^H_6_](25R)26-hydroxycholest-4-en-3-one (prepared by treatment of [25,26,26,27,27,27-^2^H_6_](25R)26-hydroxycholesterol with bacterial cholesterol oxidase), [24,24,27,27,27-^2^H_5_]3β-hydroxycholest-5-en-(25R)26-oic acid (Avanti Polar Lipids, catalog no. A80154), [24,24,27,27,27-^2^H_5_]3-oxocholest-4-en-(25R)26-oic acid (prepared by treatment of [24,24,27,27,27-^2^H_5_]3β-hydroxycholest-5-en-(25R)26-oic acid with bacterial cholesterol oxidase), 3β-hydroxycholest-5-en-(25R)26-oic acid (Avanti Polar Lipids, catalog no. A80082), 3-oxocholest-4-en-(25R)26-oic acid (Avanti Polar Lipids, catalog no. A80083), (25R)26- hydroxycholesterol (MedChemExpress, catalog no. HY-N2371), (25R)26-hydroxycholest-4-en-3-one (synthesized by Dr. Mingxing Qian and Dr. Douglas Covey, fig. S5). TO901317 was purchased from Cayman Chemical (catalog no. 71810). GSK2033 was purchased from MedChemExpress (catalog no. HY-108688). 22-NBD cholesterol was purchased from Cayman Chemical, catalog no. 30136).

### Mass spectrometry measurement of metabolites

Sputum and plasma samples from the 78 subjects with and without TB were analyzed by LC-MS(MS^3^). With minor modifications sample preparation and analysis was performed as described in^44^. In brief, plasma or sputum (2 volumes) was disinfected in methanol (8 volumes) containing [2,2,4,6,6-^2^H_5_]cholest-4-en-3-one, then 250 µL of this solution added dropwise into absolute ethanol (0.85 mL) containing isotope-labelled standards ([26,26,26,27,27,27-^2^H_6_]24R/S-hydroxycholesterol, [25,26,26,26,27,27,27-^2^H_7_]7α- hydroxycholesterol, [25,26,26,26,27,27,27-^2^H_7_]7-oxocholesterol, [25,26,26,26,27,27,27-^2^H_7_]22S- hydroxycholest-4-en-3-one, [24,24,27,27,27-^2^H_5_]3β-hydroxycholest-5-enoic acid, [27,27,27- ^2^H_3_]7α-hydroxy-3-oxocholest-4-enoic acid, [26,26,26,27,27,27-^2^H_6_]desmosterol, [25,26,26,26,27,27,27-^2^H_7_]cholesterol) under sonication and diluted to give 1.5 mL of 70% alcohol. The solution was centrifuged (17,000 x g at 4 °C for 30 min), and the supernatant applied to a C_18_ column (Sep-Pack tC_18_, 200 mg), and oxysterols and cholestenoic acids eluted in the flow-through and a 70% ethanol wash (5.5 mL). After a further wash with 70% ethanol (4 mL) cholesterol and cholest-4-en-3-one were eluted in absolute ethanol (2 mL). Both “oxysterol” and “cholesterol” fractions were divided into two equal fractions A and B.

Each fraction was lyophilized; fraction A was reconstituted in propan-2-ol (100 µL) and KH_2_PO_4_ buffer (pH 7, 1 mL) containing cholesterol oxidase (0.264 units). The mixture was incubated for 1 hr at 37 °C. The reaction was quenched by the addition of methanol (2 mL) followed by glacial acetic acid (150 µL) and [^2^H_5_]Girard P reactant (GP, 190 mg, bromide salt). The reaction was allowed to proceed overnight in the dark. Excess derivatization reagent was removed via a recycling process on an OASIS HLB column (60 mg), where the reaction mixture (3.25 mL, 69% organic) was added to the column, the eluate collected and diluted to 35% organic, recycled through the column and diluted to 17.5% organic, and recycled once more through the column. After a column wash with 10% methanol (6 mL), [^2^H_5_]GP-derivatized oxysterols were eluted in methanol (2 mL) or [^2^H_5_]GP-derivatized cholesterol in 3 mL of methanol.

Fractions B were treated in an identical fashion but in the absence of cholesterol oxidase enzyme and with the use of the [^2^H_0_]GP reagent (150 mg, chloride salt). Thus, oxysterols and cholesterol with a 3β-hydroxy-Δ^5^ function become derivatized with [^2^H_5_]GP (A fractions), while 3-oxo-Δ^4^ sterols e.g. 27-HCO, 3O-CA and cholest-4-en-3-one, become derivatized with [^2^H_0_]GP (B fractions).

LC-MS(MS^3^) analysis was performed using a Hypersil Gold C_18_ LC column (oxysterol analysis) or ACE C_18_ LC column (cholesterol analysis) linked to the electrospray ionization source of either an Orbitrap Elite or Orbitrap-IQX mass spectrometer. Quantification was performed using high resolution mass spectra (120,000 or 240,000 FWHM at *m/z* 400) against isotope-labelled standards, while MS^3^ spectra in the linear ion trap analyzer were recorded to confirm compound identification.

Samples from primary macrophage culture and the MARK-TB cohort were analyzed by the Metabolomics Core Laboratory at Washington University using liquid chromatography-tandem mass spectrometry (LC-MS/MS). Macrophages (1x10^6^) were seeded and then harvested at the desired endpoints, pelleted at 500 x g, and resuspended in 400µL methanol containing the appropriate deuterated standards. Sputum was mixed 1:1 with 0.2% dithiothreitol to liquify sputum followed by mixture with 4 parts of methanol containing the appropriate deuterated standards. Plasma was directly mixed with 4 parts methanol containing appropriate deuterated standards. All samples were incubated at room temperature in methanol for 15 minutes followed by centrifugation at 21,130 x g for 10 minutes. The supernatant was then collected for LC-MS/MS analysis. Standard curves were prepared to calculate concentrations of each oxysterol in their respective unprocessed samples (sputum, plasma, or macrophages). Samples were spiked with 8ng [2,2,4,6,6-^2^H_5_]cholesten-4-en-3-one, 16ng [25,26,26,27,27,27-^2^H_6_](25R)26-hydroxycholesterol, 16ng [25,26,26,27,27,27-^2^H_6_](25R)26-hydroxycholest-4-en-3-one, 2ng [24,24,27,27,27-^2^H_5_]3β-hydroxycholest-5-en-(25R)26-oic acid, and 2ng [24,24,27,27,27-^2^H_5_]3-oxocholest-4-en-(25R)26- oic acid. 10-point calibration curves were prepared for each oxysterol. The extracts were concentrated, reconstituted in methanol, and injected to the LC-MS/MS system for 3β-HCA, 3O- CA, cholest4-en-3-one, and 27-HC assays. The samples for 27-HC assay were derivatized with nicotinate to increase sensitivity. The 3β-HCA and 3O-CA were detected with negative multiple reaction monitoring (MRM) mode on a 4000QTRAP mass spectrometer coupled to a Shimadzu 20AD HPLC system. The cholestenone, 27-HCO, and 27-HC were detected with positive MRM mode on an API4000 mass spectrometer coupled to a Shimadzu 20AD HPLC. Data processing was conducted with Analyst 1.6.3. Calibration curves were constructed by plotting the corresponding peak area ratios of analyte/internal standard versus the corresponding analyte concentrations using weighted (1/x2) least squares regression analysis. The calibrators deviate by more than 15% of nominal concentrations were excluded from construction of calibration curve, except that deviation of 20% was acceptable for the lower limit of quantification (LLOQ), which is the low end of calibration curve. The 3β-HCA, 3O-CA, cholest-4-en-3-one, 27-HCO and 27- HC concentrations in study samples were determined from standard curves.

### Random forest classification

To assess the collective discriminative performance of the 15 oxysterols measured in this study, we trained a random forest classifier using the randomForest package (v4.7) in R.^50^ Cholesterol-normalized concentrations of the 8 discriminative oxysterols served as predictor variables, with TB status (positive or negative) as the binary outcome. To obtain stable performance estimates in the context of a moderate sample size (n = 78), we used repeated 10-fold cross-validation with 5 repeats for both model training and testing. Each forest was grown with 500 trees, with a minimum terminal node size of 3 observations to limit overfitting in small samples, and the default setting for the number of variables sampled at each split (mtry = ⌊√p⌋). For each cross-validation fold, model performance was evaluated on the held-out data by computing class probabilities for the TB-positive class and generating a receiver operating characteristic curve using the pROC package.^51^ Discriminative power was summarized as the area under the ROC curve (AUROC), averaged across folds and repetitions. Variable importance was assessed using the mean decrease in accuracy metric, calculated by permuting each predictor in the out-of-bag or validation samples and measuring the resulting loss in classification accuracy to provide an interpretable measure of each oxysterol’s contribution to TB classification within the multivariate model.

### Amplex Red Cholesterol Kit Measurement

Following extraction of the samples in methanol, cholesterol concentrations from MDM cell pellets and MARK-TB sputum samples were determined using the Amplex Red Cholesterol Assay Kit (Thermo Fisher Scientific, catalog no. A12216). The assay was performed in a 96-well optical-bottom black plate according to the manufacturer’s instructions. Following incubation at 37°C for 30 minutes, fluorescence was measured using an Agilent BioTek Synergy H1 microplate reader (excitation 530 nm and emission 590 nm). Cholesterol concentrations were calculated from a standard curve provided in the kit. Cholesterol was measured with or without preincubation with bacterial cholesterol esterase, allowing calculation of total cholesterol and free cholesterol. The esterified fraction of cholesterol was determined by subtracting free cholesterol from total cholesterol.

### Reanalysis of scRNAseq data

A publicly available scRNAseq dataset of human lung samples from 4 TB uninfected and 9 post-TB treatment patients^52^ was re-analyzed in R using the Seurat v5 package.^53^ First, the data was converted into a Seurat object then sctransform v2 was used to normalize the data and identify the 3000 most variable genes.^54^ Cells were clustered at a resolution of 1. The dataset consists of 19,632 cells that were re-annotated based on established cell lineage markers, such as *PTPRC* to define hematopoietic-derived cells, *TRAC* and *CD4* for CD4^+^ T cells, *FABP4* for alveolar macrophages, *CXCR2* and *CSF3R* for neutrophils, *KIT* for mast cells, *MAFB* and *CD14* for monocytes, *MAFB* and *SPP1* for SPP1^+^ macrophages, and *C1QA* and *MAFB* for MDMs. Gene expression was plotted with the Seurat FeaturePlot function.

### Ex vivo macrophage infections

Mtb was cultured to log-phase in Middlebrook 7H9 media with 10% OADC enrichment. A single cell suspension of bacteria was generated by low-speed centrifugation at 60 x g for 10 min. Macrophages were infected at a multiplicity of infection of 5. Plates were then centrifuged at 12 x g for 5 minutes. After 4 hours, wells were washed 3 times to remove extracellular bacteria. At 4, 48, and 72 h post-infection, cells were dissociated from wells using Accutase (Thermo Fisher Scientific, catalog no. 00-4555-56) and pelleted by centrifugation, followed by extraction with 400µL methanol containing the appropriate deuterated standards for 15 minutes at room temperature. Samples were then centrifuged at 15,000 x g for 10 minutes and the supernatant was reserved for mass spectrometry.

### LXRα reporter assay

HEK293TN cells were transduced with pGreenFire1-LXRE-in-LXRα lentiviral reporter (System Biosciences) and expanded under puromycin selection to generate a stable LXRE-luciferase reporter line, according to the manufacturer’s instructions. Cells were seeded into 96-well white plates at a density of 30,000 cells/well in complete DMEM and allowed to adhere overnight. The following day, cells were transiently transfected with an LXRα expression plasmid (LXRalpha_pcDNA3.1, Addgene plasmid #177621) and a constitutive Renilla luciferase expression plasmid as a transfection control using Lipofectamine 3000 (Invitrogen, catalog no. L3000008) in Opti-MEM reduced-serum medium, following the supplier’s protocol. After 24 h, the transfection medium was replaced with fresh complete medium containing the indicated oxysterols or 0.25% (vol/vol) DMSO vehicle control, and cells were incubated for 24 h.

Firefly luciferase and Renilla luciferase activity were then sequentially quantified using the Dual-Glo Luciferase Assay System (Promega, catalog no. E2920) according to the manufacturer’s instructions on an Agilent BioTek Synergy H1 microplate reader equipped for luminescence detection. For each experiment, background-subtracted luminescence values were normalized to the relative Renilla luciferase signal.

### Quantitative polymerase chain reaction

THP-1 cells were plated in 12-well plates at a density of 2.5 x 10^5^ per well and differentiated as described above. At the indicated time points, media was aspirated, and cells were lysed and total RNA was isolated using the RNeasy Mini Kit (Qiagen, catalog no. 74104) according to the manufacturer’s instructions. RNA concentrations were determined using a NanoDrop One spectrophotometer (Thermo Fisher Scientific), and cDNA was synthesized using the High-Capacity cDNA Reverse Transcription Kit (Thermo Fisher Scientific, catalog no. 4368814). Quantitative PCR was performed using iTaq Universal SYBR Green Supermix (BioRad, catalog no. 1725120) on a CFX Connect Real-Time System (Bio-Rad Laboratories). Fold changes in gene expression were calculated by normalizing to GAPDH as a housekeeping gene, and values are presented relative to cells treated with vehicle control. The nucleotide sequences of all primers are in table S2.

### NBD cholesterol assay

Human MDMs were infected with mCherry-expressing Δ*hsd* Mtb or the complemented strain (Δ*hsd::hsd*) at an MOI of 5. Single-cell suspensions of Mtb were prepared by low-speed centrifugation at 60 x g for 10 min.^55^ Four h post-infection, MDMs were washed to remove extracellular Mtb and resuspended in complete media containing DMSO or 1 µM LXR antagonist (GSK2033). After two days, MDMs were incubated in serum-free RPMI 1640 medium without phenol red (Gibco, Fisher Scientific, catalog no. 11835030) containing 5 µM 22NBD- Cholesterol for three hours prior to fixation with 4% paraformaldehyde for 30 minutes. Cells were stained with 4′,6-diamidino-2-phenylindole (DAPI) and mounted using Prolong Diamond Antifade (Thermo Fisher Scientific, catalog no. P36970), and images were captured using a Nikon Eclipse Ti2 confocal microscope (Nikon Instruments Inc.) equipped with 60X apochromat oil-objective lens. Image analyses were performed by selecting a region of interest around each infected macrophage and using the ROI Statistics tool in NIS-Elements version 4.40 (Nikon) to calculate the sum fluorescence intensity.

## Statistics

Analyses were conducted using GraphPad Prism version 10.3.1 for Mac (GraphPad Software, Boston, Massachusetts, USA; www.graphpad.com), or RStudio version 2023.03.1+446. The following R packages were used: ggplot2 (v3.5.1),^56^ pROC (v1.19.0.1),^51^ randomForest (v4.7),^50^ and pheatmaps (1.0.13).^57^ Normality was assessed using Shapiro-Wilks test and QQ plots. Differences between groups were assessed using 2-tailed Student’s *t* test, Wilcoxon rank sum test, Wilcoxon matched pairs signed rank test, Kruskal-Wallis test with Dunn’s post-hoc correction, 1- way ANOVA models followed by Dunnett’s multiple comparisons test, or two-way ANOVA followed by Dunnett’s multiple comparisons test, as indicated in figure legends. Analyses evaluating differences between TB groups and clinical characteristics included independent groups *t* tests, χ^2^ tests, or Fisher’s exact tests, as appropriate. A *P* value less than 0.05 was considered statistically significant.

