## Supplementary Figures S1-S5; Tables S2-S3 for "*Mycobacterium tuberculosis* inactivates host oxysterols to impair macrophage cholesterol homeostasis"

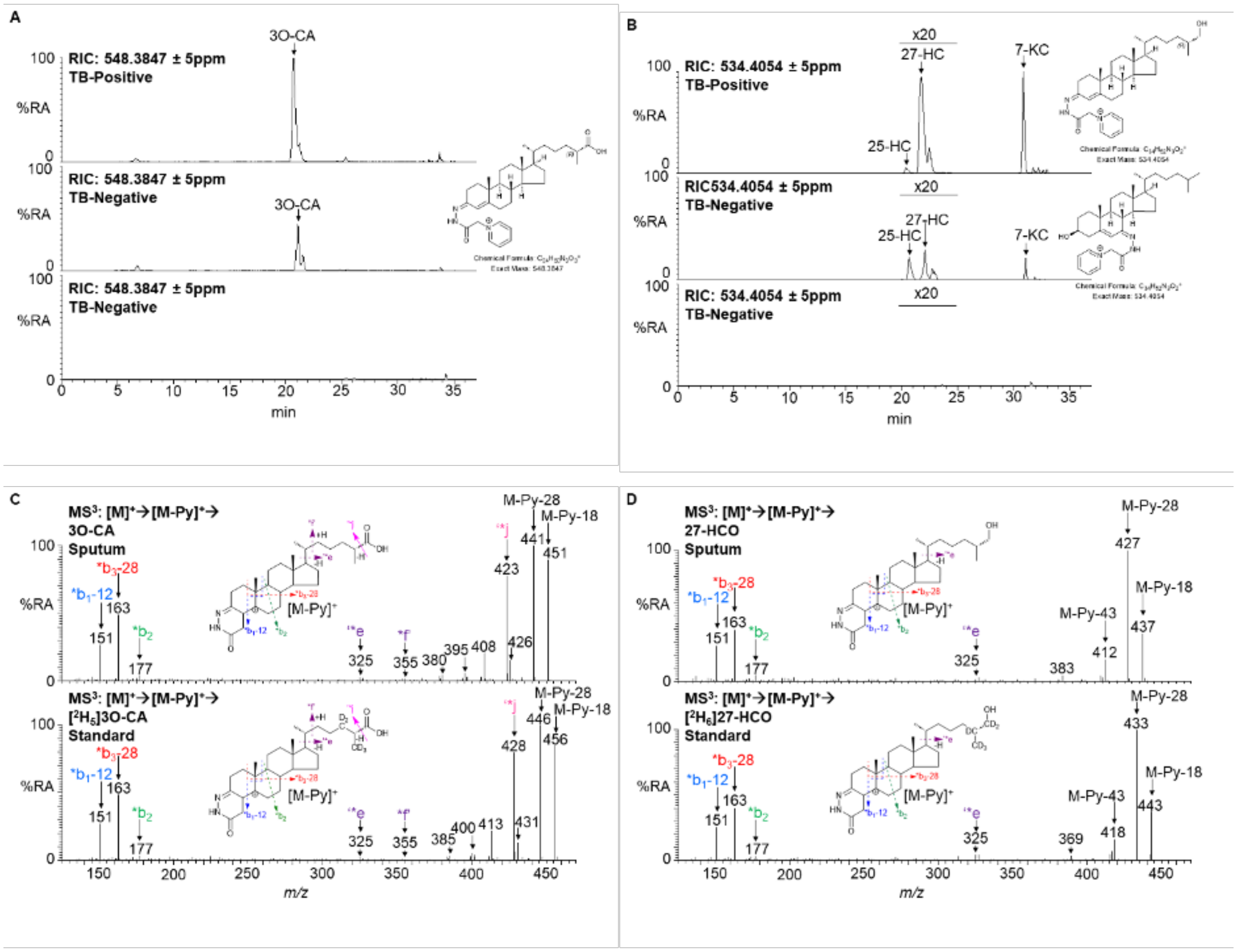


**Fig. S1. 3O-CA and 27-HCO are elevated in sputum from TB-infected individuals compared to uninfected.** Reconstructed ion chromatograms (RICs) ± 5 ppm targeting (**A**) 3O-CA and (**B**) 27-HCO, following derivatization with GP reagent. The top panels show sputum data typical of a TB-infected individual, the middle panels show data from a patient with high 3-oxo-$\Delta^4$ metabolites but not infected with TB, and the bottom panels shows data from an uninfected patient where 3-oxo-$\Delta^4$ metabolites are absent. Each chromatogram is shown on the same y-axis. GP derivatized oxysterols give cis and trans isomers about the C=N hydrazone bond which may or may not be resolved. $MS^3$ ($[M]^+ \rightarrow [M\text{-}Py]^+ \rightarrow$, Py = pyridine) of (**C**) 3O-CA from sputum and [$^2H_5$]3O-CA isotope-labelled standard, and (**D**) 27-HCO from sputum and [$^2H_6$]27-HCO isotope labelled standard. LC separation was achieved on a Hypersil Gold $C_{18}$ column utilizing a methanol, acetonitrile aqueous formic acid gradient. RICs were generated from mass spectra recorded at 240,000 resolution in the Orbitrap analyzer and $MS^3$ were recorded in the linear ion trap analyzer.

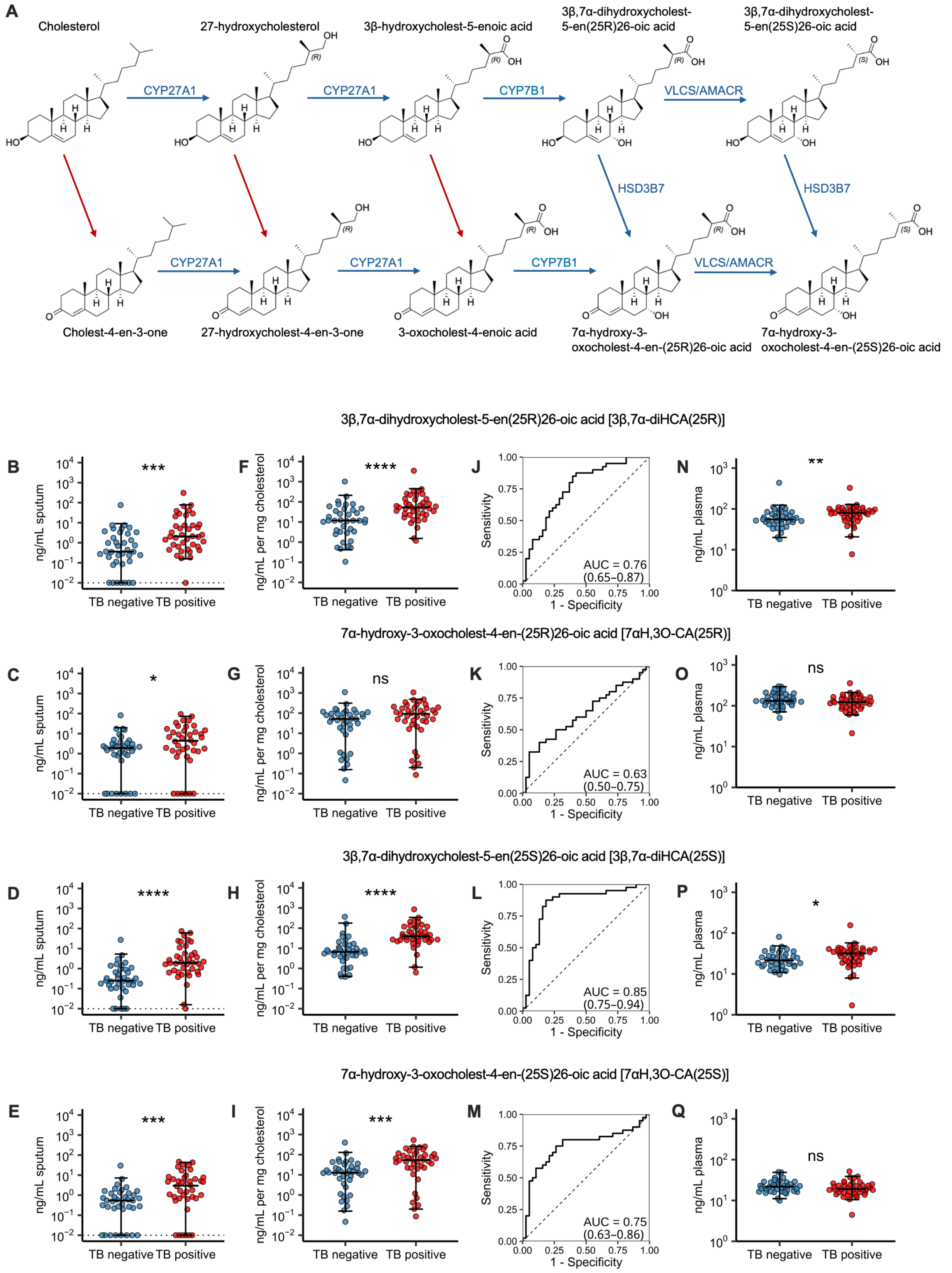


**Fig. S2. Tuberculosis is associated with accumulation of acidic pathway bile acid**

**intermediates in sputum**. **(A)** Metabolites identified in this study, with blue arrows indicating reactions in the acidic bile acid synthesis pathway, and red arrows indicating reactions that lack an established host enzyme. (**B-Q**) Sputum and plasma samples were measured using LC-MS($MS^3$). (**B-E**) Sputum concentrations, (**F-I**) cholesterol-normalized sputum concentrations, (**J-M**) ROC curves, and (**N-Q**) plasma concentrations. Statistical significance was determined using Wilcoxon rank sum tests (**B-I**; **N-Q**). Bars represent median, and whiskers encompass 95% of the observed data. The limit of detection is indicated by the dotted line. 95% confidence intervals from ROC curves estimated by the DeLong method. Abbreviations: AMACR, alpha-methyl-acyl-CoA racemase (*AMACR*); VLCS, very long chain acyl-CoA synthetase (*SLC27A2*). **** $P < 0.0001$; *** $P < 0.001$; ** $P < 0.01$; * $P < 0.05$; NS, not significant.

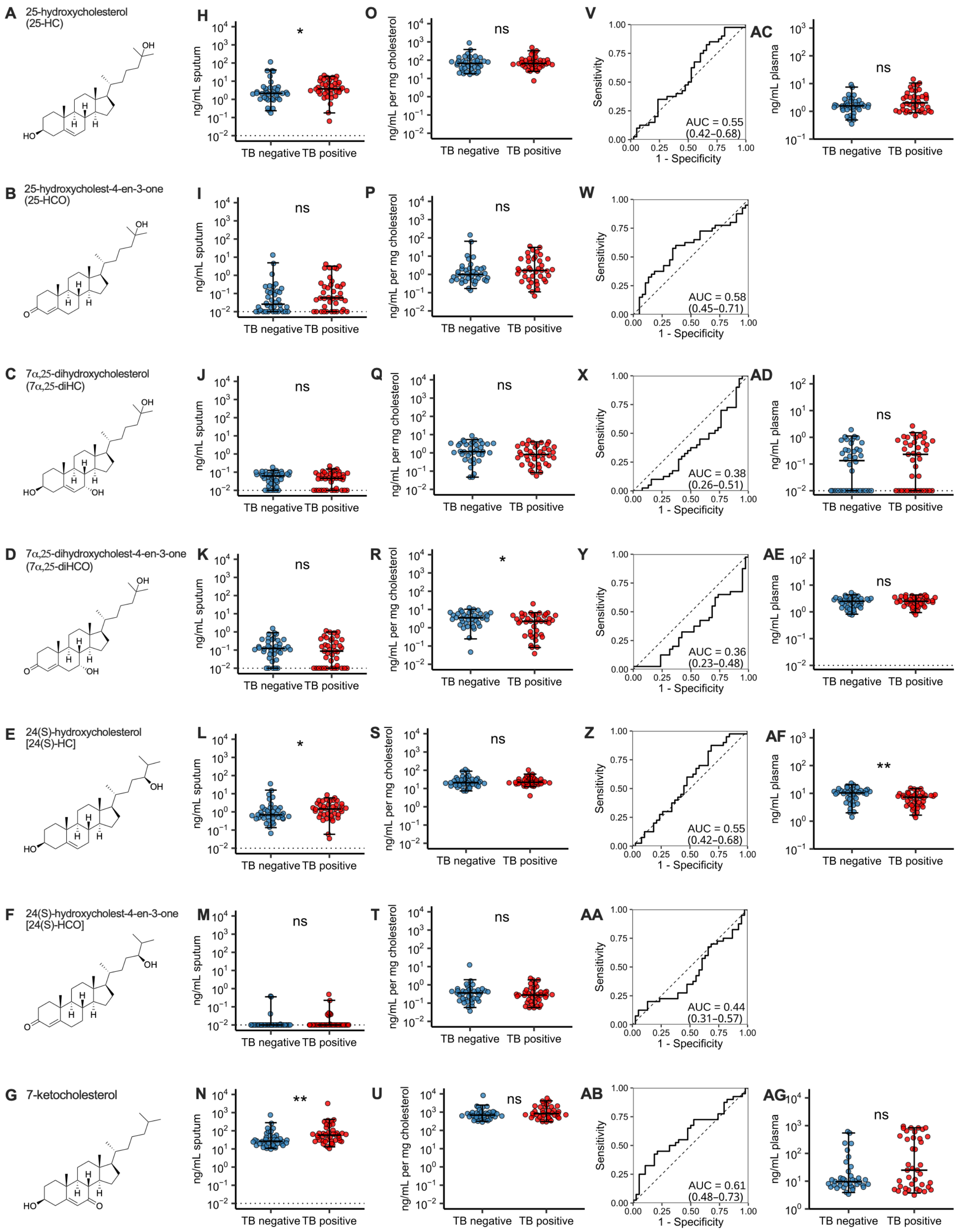


**Fig. S3. Sputum and plasma concentrations of 25-HC, 7α,25-diHC, 24(S)-HC, their 3-oxo derivatives, and 7-ketocholesterol.** Sputum and plasma oxysterols were measured using LC-

MS(MS$^3$). (**A-G**) Chemical structures, (**H-N**) sputum concentrations, (**O-U**) cholesterol-normalized sputum concentrations, (**V-AB**) ROC curves, and (**AC-AG**) plasma concentrations. Statistical significance was determined using Wilcoxon rank-sum tests (**H-U**; **AC-AG**). Bars represent median, and whiskers encompass 95% of the observed data. The limit of detection is indicated by the dotted line (**A-G**). ROC curves are shown with AUC and 95% confidence intervals estimated by the DeLong method. ** P < 0.01; * P < 0.05; NS, not significant.

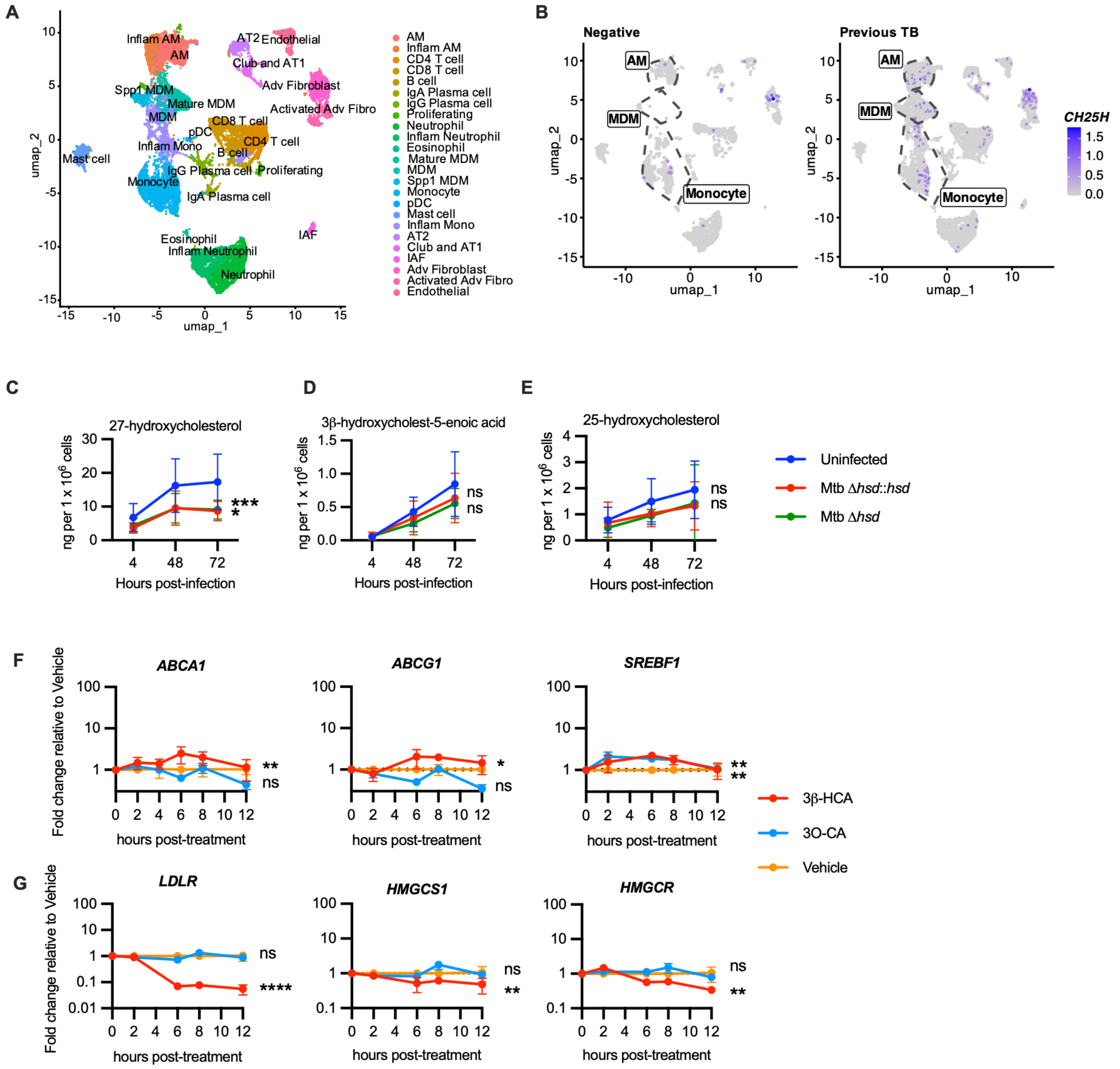


**Fig. S4. Oxysterol expression, production, and impact on gene expression.** (A) UMAP showing combined samples and cell assignments from lung resections from TB-negative individuals or those with prior history of TB. (B) Feature plots stratified by group showing expression of *CH25H*. Alveolar macrophages (AM), monocyte-derived macrophages (MDM), and monocytes are designated by dotted lines. (C-E) Human MDMs were infected with Δ*hsd* Mtb, Δ*hsd*::*hsd* Mtb, or uninfected. Cells were harvested 4, 48, and 72 hours post-infection. (C) 27-HC, (D) 3β-HCA, and (E) 25-HC concentrations were quantified by LC–MS/MS. Results are from 4 different MDM donors. (F-G) PMA-differentiated THP-1 macrophages were treated with 3β-HCA (5 µM), 3O-CA (5 µM), or vehicle control (0.25% DMSO), and cells were harvested 2, 6, 8, and 12 h after treatment. Expression is shown as fold change relative to vehicle treated controls.

Statistical significance was determined using two-way ANOVA with Dunnett's multiple comparisons test of each treatment versus uninfected (C-E) or vehicle control (F-G). **** $P < 0.0001$; *** $P<0.001$; ** $P < 0.01$; * $P < 0.05$; NS, not significant.

|

Preparation of 25(*R*)-26-hydroxycholest-4-en-3-one (3-oxo-27-hydroxycholesterol)

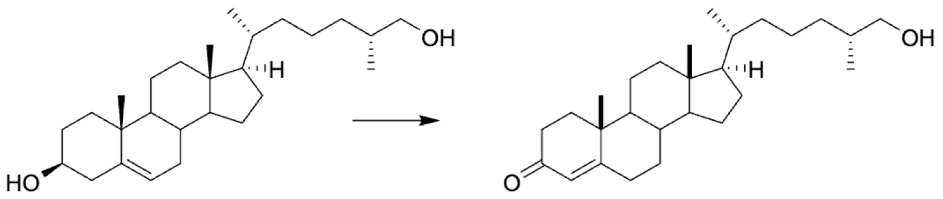

**Fig. S5. Preparation of 25(R)26-hydroxycholest-4-en-3-one.** To a solution of 25(*R*)26-hydroxycholesterol [27-hydroxycholesterol, 40 mg, 0.1 mmol) in toluene (5 mL) was added potassium bis(trimethylsilyl)amide (KHMDS, 0.5 M in toluene, 0.3 mL, 0.15 mmol) and chloro(1-phenylindeyl)bis(triphenylphosphine)ruthenium(II) (10 mg) at 23 °C. Acetone (5 mL) was added and the reaction was heated to 110 °C for 3 h. After cooling to 23 °C, aqueous $NH_4Cl$ was added and the product was extracted into EtOAc (100 mL x 2). The combined extracts were dried over anhydrous $Na_2SO_4$, filtered, solvent removed under reduced pressure and the residue was purified by flash column chromatography (silica gel, eluted with 25-35% EtOAc in hexanes) to give 3-oxo-27-hydroxycholesterol (12 mg, 30%): $^1$H NMR (400 MHz, $CDCl_3$) d 5.72 (s, 1H), 3.50-3.48 (m, 1H), 3.44-3.40 (m, 1H), 2,42-0.94 (m, 35H), 1.18 (s, 3H), 0.70 (s, 3H); $^{13}$C NMR (100 MHz, $CDCl_3$) d 199.7, 171.8, 123.7, 68.5, 55.1, 53.8, 52.8, 42.4, 39.6, 38.6, 36.1, 35.8, 35.7 (2 x C), 35.6, 34.0, 33.5, 32.9, 32.0, 28.2, 24.2, 23.4, 21.0, 18.6, 17.4, 16.5, 12.0.

**Table S2.**

Clinical and demographic characteristics of Foundation for Innovative New Diagnostics cohort.

| Variable | Peru | | P value | Vietnam | | P value |
|---|---|---|---|---|---|---|
| | TB negative (n = 19) | TB positive (n = 20) | | TB negative (n = 19) | TB positive (n = 20) | |
| Sex | | | | | | |
| Male | 9/19 (47.4%) | 6/20 (30%) | 0.265 | 7/19 (36.8%) | 5/20 (25%) | 0.423 |
| Female | 10/19 (52.6%) | 14/20 (70%) | | 12/19 (63.2%) | 15/20 (75%) | |
| Age, years; mean (SD) | 33 (11.9) | 30 (11.0) | 0.506 | 51 (16.0) | 47 (14.1) | 0.433 |
| History of BCG vaccination | 17/18 (89.5%) | 20/20 (100%) | 0.474 | 9/17 (47.4%) | 13/15 (65%) | 0.0605 |
| History of active TB | 0/7 (0%) | 1/20 (5%) | >999 | 5/19 (26.3%) | 3/20 (15%) | 0.451 |
| Normal chest x-ray | 14/16 (73.7%) | 1/19 (5%) | <0.0001 | 0/19 (0%) | 0/20 (0%) | >0.999 |
| Chest pain | 17/19 (89.5%) | 13/20 (65%) | 0.127 | 9/19 (47.4%) | 11/20 (55%) | 0.634 |
| Dyspnea | 10/19 (52.6%) | 14/20 (70%) | 0.265 | 14/19 (73.7%) | 8/20 (40%) | 0.0536 |
| Fever | 6/19 (31.6%) | 11/20 (55%) | 0.14 | 12/19 (63.2%) | 13/20 (65%) | 0.905 |
| Hemoptysis | 3/19 (15.8%) | 8/20 (40%) | 0.155 | 3/19 (15.8%) | 5/15 (25%) | 0.417 |
| Malaise | 19/19 (100%) | 19/20 (95%) | >0.999 | 16/19 (84.2%) | 14/20 (70%) | 0.451 |
| Night sweats | 3/19 (15.8%) | 12/20 (60%) | 0.0079 | 6/19 (31.6%) | 9/20 (45%) | 0.514 |
| Cough | 19/19 (100%) | 20/20 (100%) | >0.999 | 19/19 (100%) | 20/20 (100%) | >0.999 |
| Recent weight loss (without diet) | 7/19 (36.8%) | 15/20 (75%) | 0.0163 | 6/19 (31.6%) | 11/20 (55%) | 0.2 |

Reported as n (%) or mean (SD). P values calculated with $\chi 2$, Fisher's exact, or t-test, as appropriate.

**Table S3.**

qPCR primer sequences

| | |
|---|---|
| *ABCA1* | F - TTCCCGCATTATCTGGAAAGC<br>R - CAAGGTCCATTTCTTGGCTGT |
| *ABCG1* | F - ATTCAGGGACCTTTCCTATTCGG<br>R - CTCACCACTATTGAACTTCCCG |
| *GAPDH* | F - GGAGCGAGATCCCTCCAAAAT<br>R - GGCTGTTGTCATACTTCTCATGG |
| *HMGCR* | F - TGATTGACCTTTCCAGAGCAAG<br>R - CTAAAATTGCCATTCCACGAGC |
| *HMGCS1* | F - GATGTGGGAATTGTTGCCCTT<br>R - ATTGTCTCTGTTCCAACTTCCAG |
| *LDLR* | F - TACAAGTGGGTCTGCGATGG<br>R - TGAAGTCCCCGGATTTGCAG |
| *SREBF1* | F - ACAGTGACTTCCCTGGCCTAT<br>R - GCATGGACGGGTACATCTTCAA |
