## Supplementary Table S1 for "*Mycobacterium tuberculosis* inactivates host oxysterols to impair macrophage cholesterol homeostasis"

| Sputum_ID | Paired_Plasma_ID | Sex | Origin | TB Status | Smear Grad | Free cholesterol (μg/mL) | Cholest-4-en-3-one (ng/mL) | 24(S)-hydroxycholesterol (ng/mL) | 24(S)-hydroxycholest-4-en-3-one (ng/mL | 25-hydroxycholesterol (ng/mL) | 25-hydroxycholest-4-en-3-one (ng/mL) | 27-hydroxycholesterol (ng/mL) |
|---|---|---|---|---|---|---|---|---|---|---|---|---|
| S18 | P18 | Female | Peru | TB negative | 0 | 10.510 | 35.338 | 0.413 | 0.011 | 0.936 | 0.111 | 15.366 |
| S19 | P19 | Male | Peru | TB negative | 0 | 31.182 | 540.386 | 0.470 | 0.386 | 0.914 | 4.507 | 0.208 |
| S20 | P20 | Male | Peru | TB negative | 0 | 54.341 | 363.007 | 1.030 | 0.010 | 1.993 | 0.018 | 6.170 |
| S41 | P41 | Female | Peru | TB negative | 0 | 9.015 | 73.555 | 0.066 | 0.010 | 0.176 | 0.010 | 0.911 |
| S42 | P42 | Female | Peru | TB negative | 0 | 97.336 | 406.689 | 1.686 | 0.010 | 4.423 | 0.013 | 39.351 |
| S43 | P43 | Female | Peru | TB negative | 0 | 22.357 | 119.156 | 0.698 | 0.010 | 1.727 | 0.010 | 36.373 |
| S44 | P44 | Male | Peru | TB negative | 0 | 18.919 | 181.792 | 0.218 | 0.010 | 0.314 | 0.010 | 0.736 |
| S45 | P45 | Female | Peru | TB negative | 0 | 272.449 | 1463.203 | 9.043 | 0.010 | 37.656 | 0.263 | 605.105 |
| S46 | P46 | Male | Peru | TB negative | 0 | 29.592 | 178.951 | 0.314 | 0.010 | 1.092 | 0.085 | 2.346 |
| S47 | P47 | Female | Peru | TB negative | 0 | 167.398 | 879.746 | 14.706 | 0.010 | 37.564 | 1.161 | 687.146 |
| S48 | P48 | Female | Peru | TB negative | 0 | 16.235 | 128.202 | 0.417 | 0.010 | 1.944 | 0.010 | 6.467 |
| S49 | P49 | Female | Peru | TB negative | 0 | 320.603 | 341.277 | 35.390 | 0.041 | 115.653 | 0.339 | 2194.137 |
| S50 | P50 | Male | Peru | TB negative | 0 | 16.409 | 98.123 | 0.225 | 0.010 | 0.325 | 0.032 | 1.877 |
| S51 | P51 | Female | Peru | TB negative | 0 | 18.705 | 108.214 | 0.277 | 0.010 | 1.034 | 0.012 | 14.918 |
| S52 | P52 | Male | Peru | TB negative | 0 | 56.790 | 480.603 | 0.738 | 0.010 | 2.054 | 0.064 | 37.289 |
| S54 | P54 | Male | Peru | TB negative | 0 | 41.938 | 194.749 | 0.643 | 0.010 | 0.974 | 0.043 | 3.541 |
| S55 | P55 | Male | Peru | TB negative | 0 | 22.047 | 85.142 | 0.456 | 0.010 | 3.397 | 0.010 | 20.635 |
| S79 | P79 | Male | Peru | TB negative | 0 | 214.954 | 172.060 | 6.457 | 0.353 | 35.169 | 13.195 | 127.074 |
| S80 | P80 | Male | Peru | TB negative | 0 | 21.454 | 101.555 | 1.849 | 0.010 | 18.945 | 0.029 | 4.248 |
| S21 | P21 | Male | Peru | TB positive | 3 | 183.988 | 1194.501 | 4.765 | 0.011 | 18.417 | 3.181 | 14.652 |
| S22 | P22 | Male | Peru | TB positive | 3 | 260.496 | 3169.691 | 8.538 | 0.484 | 16.242 | 2.343 | 47.059 |
| S23 | P23 | Female | Peru | TB positive | 3 | 32.724 | 85.014 | 1.528 | 0.010 | 7.937 | 0.235 | 30.855 |
| S24 | P24 | Male | Peru | TB positive | 3 | 87.249 | 391.584 | 2.333 | 0.038 | 10.590 | 2.626 | 71.522 |
| S25 | P25 | Male | Peru | TB positive | 3 | 36.437 | 175.866 | 1.174 | 0.010 | 3.411 | 0.112 | 4.778 |
| S26 | P26 | Female | Peru | TB positive | 3 | 4.340 | 62.634 | 0.059 | 0.010 | 0.184 | 0.010 | 0.090 |
| S27 | P27 | Female | Peru | TB positive | 2 | 42.436 | 143.215 | 4.395 | 0.010 | 21.007 | 0.232 | 68.044 |
| S28 | P28 | Male | Peru | TB positive | 2 | 12.477 | 95.640 | 0.417 | 0.010 | 3.542 | 0.019 | 1.190 |
| S29 | P29 | Male | Peru | TB positive | 3 | 116.138 | 415.801 | 4.219 | 0.033 | 12.116 | 2.526 | 24.221 |
| S30 | P30 | Male | Peru | TB positive | 2 | 23.967 | 67.365 | 0.317 | 0.010 | 1.479 | 0.010 | 1.590 |
| S31 | P31 | Female | Peru | TB positive | 2 | 14.202 | 47.558 | 0.787 | 0.010 | 3.023 | 0.101 | 4.529 |
| S32 | P32 | Male | Peru | TB positive | 3 | 93.233 | 637.391 | 2.007 | 0.010 | 4.520 | 0.478 | 2.684 |
| S33 | P33 | Male | Peru | TB positive | 2 | 28.355 | 58.388 | 0.615 | 0.010 | 4.219 | 0.254 | 1.639 |
| S34 | P34 | Male | Peru | TB positive | 3 | 25.441 | 132.771 | 0.422 | 0.010 | 1.178 | 0.013 | 0.658 |
| S35 | P35 | Male | Peru | TB positive | 3 | 214.915 | 1166.709 | 3.909 | 0.038 | 12.772 | 2.609 | 30.033 |
| S36 | P36 | Male | Peru | TB positive | 3 | 66.748 | 435.441 | 1.598 | 0.010 | 6.536 | 0.010 | 10.106 |
| S37 | P37 | Female | Peru | TB positive | 3 | 119.158 | 718.889 | 1.663 | 0.222 | 2.995 | 4.136 | 20.854 |
| S38 | P38 | Male | Peru | TB positive | 2 | 36.786 | 69.848 | 0.723 | 0.010 | 1.963 | 0.040 | 3.985 |
| S39 | P39 | Male | Peru | TB positive | 3 | 49.356 | 122.958 | 2.988 | 0.010 | 16.048 | 0.771 | 9.737 |
| S40 | P40 | Female | Peru | TB positive | 3 | 8.530 | 7681.257 | 0.034 | 0.010 | 0.063 | 0.060 | 0.010 |
| S02 | P01 | Female | Vietnam | TB negative | 0 | 35.484 | 232.886 | 0.771 | 0.010 | 1.179 | 0.017 | 0.817 |
| S03 | P02 | Male | Vietnam | TB negative | 0 | 142.016 | 1053.662 | 5.097 | 0.010 | 2.612 | 0.243 | 6.875 |
| S04 | P03 | Male | Vietnam | TB negative | 0 | 27.587 | 214.484 | 0.561 | 0.010 | 2.411 | 0.023 | 7.551 |
| S06 | P05 | Female | Vietnam | TB negative | 0 | 16.308 | 140.795 | 0.821 | 0.010 | 2.691 | 0.010 | 8.931 |
| S07 | P06 | Male | Vietnam | TB negative | 0 | 93.801 | 299.888 | 1.349 | 0.010 | 3.922 | 0.132 | 7.903 |
| S08 | P07 | Female | Vietnam | TB negative | 0 | 24.745 | 282.270 | 1.145 | 0.010 | 2.175 | 0.018 | 20.306 |
| S09 | P08 | Male | Vietnam | TB negative | 0 | 64.175 | 439.202 | 2.657 | 0.010 | 8.972 | 0.129 | 113.592 |
| S10 | P09 | Male | Vietnam | TB negative | 0 | 32.808 | 183.441 | 0.638 | 0.010 | 2.129 | 0.116 | 30.813 |
| S11 | P11 | Male | Vietnam | TB negative | 0 | 64.302 | 343.939 | 1.667 | 0.010 | 4.820 | 0.137 | 10.044 |
| S12 | P12 | Male | Vietnam | TB negative | 0 | 21.477 | 258.359 | 0.510 | 0.010 | 1.028 | 0.045 | 8.317 |
| S13 | P13 | Male | Vietnam | TB negative | 0 | 74.622 | 402.685 | 0.948 | 0.010 | 4.214 | 0.022 | 15.800 |
| S14 | P14 | Male | Vietnam | TB negative | 0 | 23.994 | 269.749 | 0.493 | 0.010 | 1.962 | 0.010 | 1.491 |
| S15 | P15 | Female | Vietnam | TB negative | 0 | 11.628 | 138.327 | 0.144 | 0.010 | 0.242 | 0.010 | 0.992 |
| S16 | P16 | Male | Vietnam | TB negative | 0 | 83.490 | 475.865 | 1.694 | 0.010 | 4.918 | 0.190 | 12.280 |
| S17 | P17 | Male | Vietnam | TB negative | 0 | 27.696 | 278.638 | 0.622 | 0.010 | 2.320 | 0.249 | 22.953 |
| S60 | P60 | Female | Vietnam | TB negative | 0 | 37.619 | 388.684 | 1.213 | 0.010 | 2.571 | 0.019 | 15.390 |
| S61 | P61 | Female | Vietnam | TB negative | 0 | 22.901 | 188.941 | 0.674 | 0.010 | 2.136 | 0.010 | 15.485 |
| S71 | P71 | Male | Vietnam | TB negative | 0 | 24.059 | 309.219 | 0.448 | 0.010 | 1.912 | 0.010 | 3.809 |
| S72 | P72 | Female | Vietnam | TB negative | 0 | 57.728 | 435.607 | 0.427 | 0.010 | 1.282 | 0.010 | 2.234 |
| S53 | P53 | Female | Vietnam | TB positive | 3 | 64.091 | 674.983 | 1.303 | 0.010 | 3.664 | 0.052 | 5.203 |
| S56 | P56 | Male | Vietnam | TB positive | 2 | 39.300 | 405.957 | 0.700 | 0.010 | 2.770 | 0.054 | 2.546 |
| S57 | P57 | Male | Vietnam | TB positive | 3 | 84.636 | 642.862 | 2.488 | 0.010 | 8.450 | 0.129 | 34.601 |
| S58 | P58 | Male | Vietnam | TB positive | 3 | 37.612 | 488.478 | 1.465 | 0.010 | 3.293 | 0.029 | 0.723 |
| S59 | P59 | Male | Vietnam | TB positive | 2 | 25.011 | 231.539 | 0.464 | 0.010 | 1.990 | 0.010 | 1.126 |
| S62 | P62 | Female | Vietnam | TB positive | 2 | 32.364 | 70.938 | 1.050 | 0.010 | 2.081 | 0.058 | 22.595 |
| S63 | P63 | Male | Vietnam | TB positive | 2 | 47.867 | 456.177 | 1.097 | 0.010 | 1.205 | 0.010 | 5.665 |
| S64 | P64 | Male | Vietnam | TB positive | 2 | 125.149 | 728.714 | 5.892 | 0.010 | 15.280 | 0.205 | 192.359 |
| S65 | P65 | Female | Vietnam | TB positive | 2 | 18.785 | 227.276 | 0.344 | 0.010 | 0.558 | 0.010 | 1.238 |
| S66 | P66 | Male | Vietnam | TB positive | 3 | 127.886 | 894.529 | 2.306 | 0.010 | 7.274 | 0.045 | 9.017 |
| S67 | P67 | Male | Vietnam | TB positive | 3 | 34.451 | 303.447 | 0.700 | 0.010 | 3.899 | 0.056 | 22.637 |
| S68 | P68 | Male | Vietnam | TB positive | 3 | 24.756 | 227.471 | 0.505 | 0.010 | 1.362 | 0.010 | 1.281 |
| S69 | P69 | Male | Vietnam | TB positive | 2 | 152.189 | 1933.195 | 4.249 | 0.010 | 6.774 | 0.010 | 21.930 |
| S70 | P70 | Male | Vietnam | TB positive | 3 | 147.119 | 2713.756 | 1.909 | 0.010 | 5.593 | 0.301 | 0.010 |
| S73 | P73 | Female | Vietnam | TB positive | 2 | 65.524 | 384.903 | 1.357 | 0.042 | 4.036 | 0.053 | 6.134 |
| S74 | P74 | Male | Vietnam | TB positive | 3 | 178.109 | 1177.970 | 2.784 | 0.010 | 6.236 | 0.325 | 8.833 |
| S75 | P75 | Male | Vietnam | TB positive | 2 | 36.298 | 244.301 | 0.790 | 0.010 | 2.728 | 0.010 | 5.881 |
| S76 | P76 | Male | Vietnam | TB positive | 3 | 180.811 | 746.703 | 4.004 | 0.010 | 12.604 | 0.371 | 14.702 |
| S77 | P77 | Male | Vietnam | TB positive | 2 | 39.151 | 310.254 | 1.412 | 0.046 | 2.650 | 0.034 | 5.876 |
| S78 | P78 | Female | Vietnam | TB positive | 3 | 89.290 | 471.646 | 1.981 | 0.010 | 2.426 | 0.010 | 3.554 |

| Sputum_ID | 27-hydroxycholest-4-en-3-one (ng/mL) | 3β-hydroxycholest-5-enoic acid (ng/mL) | 3-oxocholest-4-enoic acid (ng/mL) | 7α,25-dihydroxycholesterol (ng/mL) | 7α,25-dihydroxycholest-4-en-3-one (ng/mL) | 3β,7α-dihydroxycholestenoic acid 25R (ng/mL) | 3-oxo,7α hydroxy cholestenoic acid 25R (ng/mL | 3β,7α-dihydroxycholestenoic acid 25S (ng/mL) |
|---|---|---|---|---|---|---|---|---|
| S18 | 3.158 | 2.569 | 0.616 | 0.010 | 0.010 | 0.010 | 0.010 | 0.010 |
| S19 | 1.740 | 0.010 | 0.695 | 0.101 | 0.036 | 0.365 | 0.967 | 0.226 |
| S20 | 1.114 | 3.729 | 0.965 | 0.062 | 0.129 | 5.709 | 15.310 | 2.961 |
| S41 | 0.010 | 0.137 | 0.010 | 0.010 | 0.010 | 0.010 | 0.010 | 0.010 |
| S42 | 0.418 | 8.930 | 0.010 | 0.080 | 0.282 | 0.242 | 2.620 | 0.326 |
| S43 | 0.164 | 11.850 | 0.664 | 0.110 | 0.090 | 0.087 | 1.769 | 0.172 |
| S44 | 0.039 | 0.151 | 0.010 | 0.020 | 0.023 | 0.010 | 0.010 | 0.010 |
| S45 | 5.825 | 40.298 | 0.522 | 0.123 | 0.390 | 3.348 | 3.733 | 0.633 |
| S46 | 0.215 | 0.599 | 0.010 | 0.023 | 0.082 | 0.348 | 3.100 | 0.268 |
| S47 | 12.327 | 120.312 | 6.772 | 0.070 | 0.869 | 7.475 | 17.426 | 1.102 |
| S48 | 0.193 | 0.575 | 0.010 | 0.084 | 0.149 | 0.010 | 1.476 | 0.109 |
| S49 | 3.278 | 199.218 | 1.315 | 0.015 | 1.557 | 3.722 | 17.309 | 0.323 |
| S50 | 0.010 | 2.074 | 0.010 | 0.047 | 0.059 | 0.194 | 1.919 | 0.160 |
| S51 | 0.150 | 2.462 | 0.010 | 0.079 | 0.059 | 0.010 | 0.831 | 0.151 |
| S52 | 0.652 | 6.024 | 0.319 | 0.040 | 0.379 | 0.745 | 2.160 | 0.255 |
| S54 | 0.327 | 0.923 | 0.010 | 0.035 | 0.116 | 0.119 | 2.401 | 0.657 |
| S55 | 0.190 | 9.308 | 0.010 | 0.074 | 0.147 | 1.034 | 3.320 | 0.100 |
| S79 | 23.925 | 31.159 | 24.240 | 0.010 | 0.010 | 7.588 | 0.010 | 1.980 |
| S80 | 0.584 | 0.010 | 2.431 | 0.185 | 0.166 | 0.010 | 0.010 | 0.010 |
| S21 | 86.727 | 23.265 | 56.243 | 0.010 | 0.414 | 23.783 | 71.598 | 27.011 |
| S22 | 82.795 | 5.567 | 176.408 | 0.217 | 0.010 | 66.574 | 45.838 | 60.117 |
| S23 | 5.564 | 2.401 | 0.521 | 0.106 | 0.010 | 0.169 | 0.705 | 0.493 |
| S24 | 31.824 | 285.137 | 15.099 | 0.127 | 0.035 | 306.531 | 93.588 | 74.840 |
| S25 | 4.315 | 3.231 | 1.782 | 0.039 | 0.067 | 2.019 | 4.077 | 2.050 |
| S26 | 0.905 | 0.010 | 2.851 | 0.010 | 0.010 | 0.349 | 1.971 | 0.519 |
| S27 | 4.636 | 6.187 | 3.792 | 0.010 | 0.872 | 1.419 | 1.772 | 0.452 |
| S28 | 0.776 | 0.010 | 0.713 | 0.046 | 0.026 | 0.445 | 1.196 | 0.636 |
| S29 | 96.399 | 48.315 | 45.329 | 0.010 | 0.010 | 41.500 | 0.010 | 24.353 |
| S30 | 2.297 | 1.132 | 0.960 | 0.010 | 0.010 | 2.101 | 0.010 | 1.638 |
| S31 | 3.009 | 1.073 | 0.936 | 0.010 | 0.010 | 0.640 | 0.010 | 0.379 |
| S32 | 17.213 | 2.530 | 4.214 | 0.010 | 0.010 | 2.488 | 1.382 | 1.639 |
| S33 | 0.836 | 1.295 | 2.105 | 0.113 | 0.010 | 1.507 | 0.459 | 1.055 |
| S34 | 0.475 | 0.136 | 0.505 | 0.072 | 0.055 | 0.224 | 0.693 | 0.016 |
| S35 | 59.088 | 39.752 | 35.534 | 0.047 | 0.131 | 74.770 | 44.989 | 44.508 |
| S36 | 7.226 | 7.875 | 2.136 | 0.072 | 0.171 | 28.527 | 17.102 | 22.239 |
| S37 | 27.032 | 0.010 | 23.070 | 0.010 | 0.010 | 0.183 | 5.968 | 3.080 |
| S38 | 0.251 | 0.756 | 0.611 | 0.074 | 0.014 | 0.468 | 1.624 | 0.815 |
| S39 | 14.139 | 3.055 | 1.968 | 0.010 | 0.010 | 3.306 | 0.010 | 1.904 |
| S40 | 5.043 | 0.010 | 3.227 | 0.010 | 0.010 | 0.010 | 0.010 | 0.010 |
| S02 | 0.113 | 0.206 | 0.010 | 0.108 | 0.010 | 0.978 | 0.010 | 0.179 |
| S03 | 0.091 | 1.279 | 0.010 | 0.010 | 0.442 | 1.340 | 2.330 | 0.878 |
| S04 | 0.223 | 6.058 | 0.010 | 0.010 | 0.341 | 5.146 | 1.856 | 4.664 |
| S06 | 0.058 | 2.282 | 0.010 | 0.075 | 0.029 | 0.977 | 1.811 | 0.342 |
| S07 | 0.234 | 1.536 | 0.010 | 0.095 | 0.154 | 0.010 | 0.803 | 0.036 |
| S08 | 0.239 | 1.633 | 0.010 | 0.028 | 0.084 | 0.443 | 1.348 | 0.010 |
| S09 | 0.190 | 37.583 | 0.010 | 0.017 | 0.441 | 5.033 | 9.095 | 1.678 |
| S10 | 2.010 | 5.169 | 0.752 | 0.055 | 0.070 | 0.111 | 2.974 | 0.179 |
| S11 | 0.163 | 1.041 | 0.010 | 0.010 | 0.382 | 0.292 | 1.900 | 0.496 |
| S12 | 0.015 | 1.479 | 0.010 | 0.070 | 0.161 | 0.239 | 1.071 | 0.240 |
| S13 | 0.010 | 54.903 | 0.794 | 0.109 | 0.761 | 75.899 | 81.493 | 27.263 |
| S14 | 0.010 | 0.694 | 0.010 | 0.077 | 0.117 | 0.078 | 0.453 | 0.104 |
| S15 | 0.031 | 0.104 | 0.010 | 0.039 | 0.010 | 0.273 | 0.010 | 0.073 |
| S16 | 0.509 | 3.806 | 0.010 | 0.030 | 0.758 | 3.313 | 6.876 | 1.398 |
| S17 | 2.539 | 11.028 | 0.991 | 0.106 | 0.107 | 0.698 | 4.541 | 0.152 |
| S60 | 0.401 | 4.216 | 0.010 | 0.102 | 0.211 | 2.835 | 4.939 | 1.449 |
| S61 | 0.139 | 1.585 | 0.010 | 0.045 | 0.133 | 0.234 | 2.171 | 0.330 |
| S71 | 0.100 | 1.496 | 0.010 | 0.062 | 0.106 | 1.753 | 0.899 | 0.937 |
| S72 | 0.010 | 0.240 | 0.010 | 0.070 | 0.105 | 0.357 | 0.010 | 0.224 |
| S53 | 14.752 | 6.569 | 16.357 | 0.010 | 0.348 | 5.764 | 22.322 | 2.701 |
| S56 | 6.410 | 3.667 | 3.584 | 0.045 | 0.160 | 2.456 | 6.737 | 1.804 |
| S57 | 13.128 | 22.847 | 9.382 | 0.045 | 0.372 | 20.642 | 11.453 | 11.395 |
| S58 | 9.474 | 2.800 | 9.917 | 0.139 | 0.192 | 5.198 | 9.306 | 2.225 |
| S59 | 4.189 | 0.670 | 3.014 | 0.053 | 0.079 | 0.421 | 3.295 | 0.655 |
| S62 | 1.338 | 1.481 | 0.596 | 0.094 | 0.010 | 0.369 | 0.010 | 0.154 |
| S63 | 6.665 | 2.408 | 9.346 | 0.073 | 0.087 | 6.716 | 8.760 | 5.723 |
| S64 | 5.256 | 40.610 | 4.600 | 0.082 | 0.596 | 7.367 | 24.014 | 3.390 |
| S65 | 0.917 | 0.332 | 1.112 | 0.092 | 0.039 | 0.646 | 1.937 | 0.654 |
| S66 | 19.334 | 13.207 | 9.006 | 0.069 | 0.608 | 28.314 | 34.680 | 13.679 |
| S67 | 2.149 | 6.872 | 2.913 | 0.088 | 0.107 | 0.798 | 1.283 | 0.704 |
| S68 | 2.019 | 3.326 | 3.954 | 0.042 | 0.156 | 1.155 | 11.861 | 1.794 |
| S69 | 7.610 | 8.490 | 5.980 | 0.070 | 0.482 | 8.200 | 13.619 | 6.189 |
| S70 | 83.060 | 12.820 | 47.213 | 0.098 | 0.963 | 1.970 | 14.523 | 3.708 |
| S73 | 3.928 | 4.323 | 1.679 | 0.013 | 0.426 | 4.790 | 4.642 | 4.104 |
| S74 | 23.919 | 20.147 | 15.958 | 0.148 | 0.987 | 38.727 | 24.904 | 21.955 |
| S75 | 4.289 | 2.260 | 3.738 | 0.029 | 0.171 | 0.814 | 1.430 | 0.985 |
| S76 | 10.235 | 11.112 | 10.417 | 0.043 | 1.146 | 7.330 | 9.022 | 4.739 |
| S77 | 1.760 | 1.787 | 3.441 | 0.010 | 0.088 | 1.013 | 2.549 | 1.136 |
| S78 | 2.362 | 1.485 | 6.414 | 0.043 | 0.267 | 1.777 | 6.395 | 1.989 |

| Sputum_ID | 3-oxo,7α hydroxy cholestenoic acid 25S (ng/mL) | 7-ketocholesterol (ng/mL) | Age (Years) | History of BCG vaccination | History of active TB | Normal chest x-ray | Chest pain | Dyspnea | Fever | Hemoptysis | Malaise | Night sweats | Cough | Weight Loss |
|---|---|---|---|---|---|---|---|---|---|---|---|---|---|---|
| S18 | 0.010 | 19.870 | 39.3 | Yes | No | Yes | Yes | Yes | No | Yes | Yes | No | Yes | No |
| S19 | 0.261 | 256.877 | 30.3 | Yes | No | Yes | Yes | No | Yes | No | Yes | No | Yes | No |
| S20 | 6.420 | 34.034 | 41.3 | Yes | No | Yes | Yes | Yes | No | Yes | Yes | No | Yes | Yes |
| S41 | 0.010 | 12.103 | 26.4 | No | Unknown | Yes | Yes | No | No | No | Yes | No | Yes | Yes |
| S42 | 0.540 | 30.859 | 51.4 | Yes | Unknown | No | Yes | No | No | No | Yes | Yes | Yes | No |
| S43 | 0.220 | 15.369 | 23.4 | Yes | Unknown | Yes | Yes | No | Yes | No | Yes | No | Yes | Yes |
| S44 | 0.010 | 15.685 | 23.5 | Yes | Unknown | Yes | Yes | No | No | No | Yes | No | Yes | No |
| S45 | 0.979 | 94.162 | 51.5 | Yes | Unknown | Yes | Yes | Yes | No | No | Yes | Yes | Yes | Yes |
| S46 | 0.494 | 30.508 | 20.5 | Yes | Unknown | Yes | Yes | Yes | Yes | No | Yes | No | Yes | Yes |
| S47 | 2.477 | 135.829 | 20.6 | Yes | Unknown | Yes | Yes | Yes | Yes | No | Yes | No | Yes | No |
| S48 | 0.200 | 11.533 | 39.5 | Yes | Unknown | Yes | Yes | No | No | No | Yes | No | Yes | Yes |
| S49 | 2.089 | 91.677 | 24.5 | Yes | Unknown | Yes | Yes | Yes | No | No | Yes | No | Yes | No |
| S50 | 0.278 | 9.710 | 29.7 | Yes | Unknown | Yes | No | No | No | No | Yes | No | Yes | No |
| S51 | 0.269 | 16.347 | 46.6 | Yes | Unknown | No | Yes | Yes | No | Yes | Yes | No | Yes | No |
| S52 | 0.701 | 19.508 | 55.7 | Yes | Unknown | No | No | No | No | No | Yes | No | Yes | No |
| S54 | 1.357 | 32.351 | 20.0 | Yes | No | Unknown | Yes | No | No | No | Yes | No | Yes | Yes |
| S55 | 0.793 | 21.098 | 35.0 | Unknown | No | Unknown | Yes | Yes | Yes | No | Yes | Yes | Yes | No |
| S79 | 0.010 | 138.106 | 24.1 | Yes | No | Yes | Yes | Yes | Yes | No | Yes | No | Yes | No |
| S80 | 0.010 | 741.996 | 22.1 | Yes | No | Unknown | Yes | Yes | No | No | Yes | No | Yes | No |
| S21 | 43.066 | 400.072 | 41.7 | Yes | No | Unknown | No | Yes | Yes | Yes | Yes | Yes | Yes | Yes |
| S22 | 30.310 | 315.536 | 21.6 | Yes | No | No | No | Yes | No | No | Yes | No | Yes | No |
| S23 | 0.193 | 70.773 | 21.5 | Yes | No | No | Yes | Yes | Yes | Yes | Yes | Yes | Yes | No |
| S24 | 46.584 | 33.864 | 19.5 | Yes | No | No | Yes | Yes | No | No | Yes | No | Yes | Yes |
| S25 | 4.915 | 72.212 | 26.4 | Yes | No | No | No | No | No | No | Yes | No | Yes | Yes |
| S26 | 1.086 | 13.087 | 22.4 | Yes | No | No | Yes | Yes | Yes | No | Yes | Yes | Yes | No |
| S27 | 1.454 | 3182.508 | 28.4 | Yes | No | No | No | Yes | No | Yes | Yes | No | Yes | Yes |
| S28 | 0.685 | 327.659 | 40.3 | Yes | No | No | No | No | No | No | Yes | No | Yes | Yes |
| S29 | 0.010 | 229.873 | 21.1 | Yes | No | No | Yes | No | Yes | No | Yes | Yes | Yes | Yes |
| S30 | 0.010 | 30.187 | 43.4 | Yes | No | No | Yes | No | Yes | No | No | Yes | Yes | Yes |
| S31 | 0.010 | 150.011 | 19.4 | Yes | No | No | Yes | Yes | Yes | Yes | Yes | Yes | Yes | Yes |
| S32 | 0.995 | 371.890 | 25.4 | Yes | Yes | No | Yes | Yes | No | Yes | Yes | Yes | Yes | Yes |
| S33 | 0.641 | 25.712 | 26.4 | Yes | No | No | Yes | Yes | Yes | Yes | Yes | No | Yes | No |
| S34 | 0.530 | 30.195 | 38.5 | Yes | No | No | Yes | Yes | No | No | Yes | No | Yes | Yes |
| S35 | 33.614 | 226.321 | 22.6 | Yes | No | No | Yes | Yes | No | Yes | Yes | No | Yes | No |
| S36 | 16.885 | 74.590 | 35.6 | Yes | No | No | Yes | Yes | Yes | Yes | Yes | Yes | Yes | Yes |
| S37 | 4.282 | 79.224 | 52.3 | Yes | No | No | Yes | Yes | Yes | No | Yes | Yes | Yes | Yes |
| S38 | 1.014 | 26.759 | 53.3 | Yes | No | Yes | No | No | Yes | No | Yes | Yes | Yes | Yes |
| S39 | 0.010 | 197.845 | 30.1 | Yes | No | No | Yes | No | No | No | Yes | Yes | Yes | Yes |
| S40 | 0.010 | 49.320 | 19.3 | Yes | No | No | No | Yes | Yes | No | Yes | Yes | Yes | Yes |
| S02 | 0.010 | 72.457 | 53.4 | No | Yes | No | No | Yes | No | No | Yes | No | Yes | No |
| S03 | 1.203 | 53.038 | 29.5 | Yes | Yes | No | Yes | Yes | Yes | Yes | Yes | No | Yes | No |
| S04 | 1.601 | 26.368 | 59.4 | Yes | No | No | Yes | Yes | Yes | No | Yes | Yes | Yes | Yes |
| S06 | 0.723 | 14.605 | 48.4 | Yes | No | No | No | Yes | Yes | No | Yes | Yes | Yes | No |
| S07 | 0.228 | 57.365 | 73.4 | No | No | No | Yes | Yes | Yes | No | No | Yes | Yes | No |
| S08 | 0.539 | 46.078 | 70.4 | Unknown | No | No | No | Yes | Yes | No | Yes | No | Yes | No |
| S09 | 3.409 | 35.868 | 36.3 | Yes | No | No | No | No | Yes | No | Yes | Yes | Yes | Yes |
| S10 | 0.606 | 18.751 | 26.6 | Yes | No | No | No | No | No | No | No | No | Yes | No |
| S11 | 0.792 | 25.629 | 49.5 | Yes | No | No | Yes | Yes | Yes | No | Yes | No | Yes | Yes |
| S12 | 0.264 | 17.127 | 45.4 | Yes | No | No | Yes | No | Yes | No | Yes | No | Yes | No |
| S13 | 29.836 | 30.417 | 59.3 | No | No | No | Yes | Yes | Yes | Yes | Yes | Yes | Yes | Yes |
| S14 | 0.312 | 10.819 | 69.4 | No | Yes | No | Yes | No | No | Yes | Yes | No | Yes | No |
| S15 | 0.010 | 24.715 | 30.5 | No | No | No | No | Yes | No | No | Yes | No | Yes | No |
| S16 | 1.810 | 43.527 | 46.3 | Yes | No | No | Yes | Yes | Yes | No | Yes | Yes | Yes | No |
| S17 | 1.243 | 27.270 | 35.5 | No | No | No | No | Yes | No | No | Yes | No | Yes | No |
| S60 | 1.620 | 21.202 | 32.5 | No | No | No | No | Yes | No | No | Yes | No | Yes | No |
| S61 | 0.596 | 24.350 | 76.5 | Yes | No | No | No | Yes | No | No | Yes | No | Yes | No |
| S71 | 0.363 | 15.491 | 63.5 | Unknown | Yes | No | No | No | Yes | No | No | No | Yes | Yes |
| S72 | 0.010 | 30.382 | 61.5 | No | Yes | No | Yes | Yes | Yes | No | Yes | No | Yes | Yes |
| S53 | 6.250 | 43.721 | 30.4 | Yes | No | No | Yes | No | Yes | No | Yes | No | Yes | No |
| S56 | 3.199 | 28.141 | 33.9 | Yes | No | No | No | Yes | Yes | No | Yes | Yes | Yes | Yes |
| S57 | 7.547 | 39.673 | 49.8 | Yes | No | No | Yes | Yes | Yes | No | Yes | Yes | Yes | Yes |
| S58 | 2.765 | 27.040 | 56.8 | Unknown | No | No | Yes | No | No | No | Yes | No | Yes | No |
| S59 | 1.650 | 10.496 | 47.4 | Unknown | No | No | No | No | No | No | No | No | Yes | No |
| S62 | 0.010 | 61.444 | 60.5 | No | No | No | No | No | Yes | No | Yes | Yes | Yes | No |
| S63 | 5.077 | 25.795 | 33.4 | Yes | No | No | No | Yes | Yes | No | Yes | Yes | Yes | Yes |
| S64 | 7.634 | 106.493 | 35.5 | Yes | No | No | Yes | Yes | No | No | Yes | No | Yes | Yes |
| S65 | 0.825 | 17.210 | 56.5 | Yes | No | No | Yes | Yes | Yes | Yes | Yes | Yes | Yes | Yes |
| S66 | 26.256 | 63.161 | 56.5 | Yes | No | No | Yes | Yes | Yes | Yes | Yes | Yes | Yes | Yes |
| S67 | 0.689 | 18.830 | 24.5 | Yes | No | No | No | No | Yes | Yes | Yes | Yes | Yes | Yes |
| S68 | 4.611 | 64.573 | 46.4 | Yes | No | No | Yes | No | Yes | Yes | Yes | Yes | Yes | Yes |
| S69 | 6.385 | 72.912 | 57.3 | Yes | No | No | Yes | No | Yes | No | Yes | Yes | Yes | Yes |
| S70 | 9.597 | 44.654 | 60.5 | Unknown | No | No | No | No | No | No | No | No | Yes | No |
| S73 | 4.848 | 54.124 | 60.4 | Unknown | Yes | No | No | No | Yes | No | No | No | Yes | No |
| S74 | 19.132 | 52.464 | 69.4 | Yes | No | No | No | No | No | No | No | No | Yes | No |
| S75 | 0.814 | 25.270 | 59.5 | Yes | No | No | Yes | Yes | Yes | No | Yes | No | Yes | No |
| S76 | 8.177 | 61.038 | 48.9 | No | Yes | No | Yes | No | No | No | No | No | Yes | No |
| S77 | 1.789 | 19.923 | 33.8 | Yes | Yes | No | Yes | Yes | Yes | Yes | Yes | No | Yes | Yes |
| S78 | 4.691 | 66.176 | 20.4 | Unknown | No | No | No | No | No | No | No | No | Yes | Yes |

| Plasma_ID | Paired_Sputum _I | Smear Grad | TB Status | Origin | Sex | Free cholesterol (µg/mL) | Cholest-4-en-3-one (ng/mL) | 24(S)-hydroxycholesterol (ng/mL) | 25-hydroxycholesterol (ng/mL) | 27-hydroxycholesterol (ng/mL) | 27-hydroxycholest-4-en-3-one (ng/mL) | 3β-hydroxycholest-5-enoic acid (ng/mL) | 3-oxocholest-4-enoic acid (ng/mL) | 7α,25-dihydroxycholesterol (ng/mL) | 7α,25-dihydroxycholest-4-en-3-one (ng/mL) | 7α,26-dihydroxycholesterol (ng/mL) | 7α,26-dihydroxycholest-4-en-3-one (ng/m | 3β,7α-dihydroxycholest-5-en-(25R)26-oic acid | 7α-hydroxy-3-oxocholest-4-en-(25R)26-oic acid ( | 3β,7α-dihydroxycholest-5-en-(25S)26-oic acid (ng/mL) | 7α-hydroxy-3-oxocholest-4-en-(25S)26-oic acid (ng/m | 7-ketocholesterol (ng/mL) |
|---|---|---|---|---|---|---|---|---|---|---|---|---|---|---|---|---|---|---|---|---|---|---|
| P01 | S19 | 0 | TB negative | Peru | Male | 255.2 | 84.75 | 11.6 | 2.5 | 25.32 | 0.52 | 226.8 | 2.25 | 0.15 | 1.415 | 0.475 | 4.865 | 100.35 | 122.4 | 36.4 | 22.15 | 75 |
| P02 | S20 | 0 | TB negative | Peru | Male | 312.15 | 106.15 | 13.9 | 9.2 | 33.735 | 0.545 | 179.15 | 2.25 | 0.835 | 0.765 | 3.19 | 2.565 | 67.55 | 109.95 | 17.55 | 15.85 | 611.5 |
| P03 | S18 | 0 | TB negative | Peru | Female | 302.8 | 80.15 | 7.7 | 3.15 | 31.03 | 0.785 | 149.2 | 3.05 | 0.195 | 1.91 | 0.5 | 8.485 | 436.8 | 293 | 80.8 | 48.65 | 228 |
| P05 | S08 | 0 | TB negative | Vietnam | Female | 310.6 | 57 | 13.35 | 1.55 | 15.125 | 0.575 | 75.9 | 2.8 | 0.82 | 2.84 | 2.29 | 9.955 | 25.65 | 210.35 | 12.35 | 36.1 | 22.35 |
| P06 | S03 | 0 | TB negative | Vietnam | Male | 281.55 | 137.15 | 12.7 | 1.25 | 16.21 | 0.595 | 103.85 | 2.9 | 0.01 | 2.715 | 0.01 | 8.915 | 74.05 | 182.9 | 29.9 | 27.75 | 5.7 |
| P07 | S09 | 0 | TB negative | Vietnam | Male | 357.45 | 111.95 | 6.65 | 1.65 | 13.67 | 0.395 | 123.5 | 3.2 | 0.01 | 3.91 | 0.01 | 11.09 | 62.15 | 200.5 | 38.15 | 28.55 | 7.65 |
| P08 | S11 | 0 | TB negative | Vietnam | Male | 319.9 | 67.3 | 6.95 | 1.7 | 39.96 | 0.4 | 133.95 | 2.4 | 0.175 | 2.705 | 1.93 | 14.385 | 43.3 | 210.55 | 13.45 | 28.85 | 10.3 |
| P09 | S15 | 0 | TB negative | Vietnam | Female | 329.15 | 114.15 | 15 | 1.35 | 26.425 | 0.455 | 109.35 | 2.45 | 0.01 | 3.69 | 0.01 | 12.645 | 33.8 | 166.65 | 15.2 | 22.35 | 6.15 |
| P11 | S07 | 0 | TB negative | Vietnam | Male | 518.2 | 115.65 | 12.7 | 1.85 | 34.66 | 0.535 | 156.15 | 1.4 | 0.36 | 2.48 | 0.71 | 8.59 | 78.95 | 90.9 | 24.35 | 16.35 | 9.35 |
| P12 | S13 | 0 | TB negative | Vietnam | Male | 349.9 | 120.1 | 9.25 | 1.1 | 17.675 | 0.42 | 86.05 | 3.4 | 0.01 | 4.145 | 0.01 | 18.05 | 47.8 | 250.55 | 18.4 | 32 | 6.45 |
| P13 | S16 | 0 | TB negative | Vietnam | Male | 405.1 | 182.15 | 11.7 | 1.6 | 49.915 | 0.33 | 210.3 | 3.25 | 0.01 | 3.4 | 0.01 | 9.73 | 111.6 | 161.25 | 46.55 | 22.55 | 8.45 |
| P14 | S12 | 0 | TB negative | Vietnam | Male | 458.95 | 98.05 | 10.3 | 1.75 | 22.825 | 0.535 | 138.2 | 2.2 | 0.295 | 1.875 | 0.885 | 6.225 | 48.6 | 104.85 | 18.65 | 16.15 | 6.85 |
| P15 | S10 | 0 | TB negative | Vietnam | Male | 323.8 | 96.5 | 4.05 | 1.55 | 23.44 | 0.405 | 158.95 | 2.55 | 0.01 | 2.375 | 0.01 | 10.14 | 60.3 | 159.95 | 26.75 | 22.95 | 5.9 |
| P16 | S04 | 0 | TB negative | Vietnam | Male | 365.55 | 95.35 | 20.2 | 2.05 | 28.685 | 0.295 | 173.55 | 2.1 | 0.01 | 3.175 | 0.01 | 11.89 | 110.15 | 130.95 | 44.25 | 17.85 | 9.35 |
| P17 | S06 | 0 | TB negative | Vietnam | Female | 466.45 | 108.3 | 16.35 | 2.75 | 34.68 | 0.29 | 149.35 | 1.15 | 0.96 | 0.955 | 4.18 | 4.38 | 59.1 | 123.45 | 17.65 | 17.9 | 13 |
| P18 | S17 | 0 | TB negative | Vietnam | Male | 406 | 87.55 | 18.4 | 2.25 | 25.13 | 0.605 | 171.3 | 2.4 | 1.045 | 1.74 | 4.955 | 7.31 | 60.45 | 201.8 | 16 | 27.9 | 11.2 |
| P19 | S02 | 0 | TB negative | Vietnam | Female | 542.35 | 155.75 | 10.45 | 1.35 | 26.845 | 0.67 | 95.7 | 2.35 | 0.095 | 2.900 | 0.54 | 8.035 | 20.2 | 142.4 | 9.7 | 21.4 | 7.1 |
| P20 | S14 | 0 | TB negative | Vietnam | Male | 400.8 | 72.7 | 6.2 | 7.35 | 14.835 | 0.39 | 88.6 | 1.65 | 1.065 | 1.64 | 2.53 | 4.37 | 36.5 | 131.5 | 12.1 | 23 | 539.7 |
| P21 | S21 | 3 | TB positive | Peru | Male | 318.75 | 163.35 | 6.55 | 3.1 | 36.945 | 0.79 | 212.5 | 1.95 | 2.645 | 2.715 | 9.33 | 9.21 | 109.55 | 190.15 | 41.5 | 38.5 | 253.95 |
| P22 | S36 | 3 | TB positive | Peru | Male | 387 | 146.3 | 1.65 | 0.7 | 9.545 | 0.525 | 180.3 | 2.5 | 1.13 | 3.205 | 2.26 | 6.37 | 96.75 | 163.25 | 30.65 | 26.15 | 24.85 |
| P23 | S35 | 3 | TB positive | Peru | Male | 235.9 | 92.5 | 9.75 | 5.45 | 3.845 | 0.235 | 132.5 | 1.05 | 0.035 | 2.08 | 0.01 | 4.525 | 92.7 | 60.9 | 43.85 | 12.25 | 425.05 |
| P24 | S40 | 3 | TB positive | Peru | Female | 382.45 | 104.05 | 13.4 | 14.4 | 13.65 | 0.405 | 141.85 | 1.9 | 1.465 | 1.99 | 3.15 | 4.91 | 78.5 | 114.6 | 32.05 | 23.8 | 946.3 |
| P25 | S39 | 3 | TB positive | Peru | Male | 270.3 | 144.55 | 3.55 | 7.5 | 31.935 | 0.38 | 200.3 | 2.25 | 0.14 | 2.32 | 1.12 | 6.38 | 91.05 | 78.8 | 41.8 | 14.1 | 829.1 |
| P26 | S38 | 2 | TB positive | Peru | Male | 249.15 | 175.9 | 8.7 | 3.3 | 5.565 | 0.43 | 222.1 | 3.05 | 0.01 | 3.385 | 0.01 | 12.145 | 91.6 | 132.55 | 37.5 | 20.3 | 145.2 |
| P27 | S33 | 2 | TB positive | Peru | Male | 238.9 | 78.75 | 2.5 | 4 | 13.565 | 0.665 | 174.55 | 2.65 | 0.23 | 2.275 | 0.405 | 6.34 | 89.05 | 124.25 | 40.75 | 22.3 | 602.6 |
| P28 | S29 | 3 | TB positive | Peru | Male | 236.75 | 181.85 | 5 | 5.8 | 1.165 | 0.485 | 216.8 | 3.4 | 0.01 | 1.77 | 0.01 | 4.505 | 112.4 | 64.15 | 48.2 | 11.95 | 647.25 |
| P29 | S30 | 2 | TB positive | Peru | Male | 248 | 162.9 | 1.35 | 1.05 | 12.765 | 0.375 | 149.9 | 1.8 | 0.01 | 2.46 | 0.01 | 7.2 | 79.7 | 70.05 | 29.4 | 13.1 | 38.4 |
| P30 | S31 | 2 | TB positive | Peru | Female | 111.1 | 30.1 | 3.5 | 3.4 | 3.495 | 0.26 | 30 | 0.55 | 0.335 | 0.95 | 1.04 | 1.01 | 7.75 | 21.25 | 1.7 | 4.45 | 321.45 |
| P31 | S32 | 3 | TB positive | Peru | Male | 298.45 | 89.25 | 3 | 4.55 | 9.695 | 0.69 | 149.1 | 2.2 | 0.23 | 1.255 | 0.75 | 4 | 59.05 | 95.1 | 19.65 | 15.95 | 401.25 |
| P32 | S24 | 3 | TB positive | Peru | Male | 183.25 | 92.25 | 2.9 | 6.05 | 8.13 | 0.315 | 213.55 | 2.4 | 0.01 | 3.57 | 0.01 | 7.635 | 106.4 | 93.25 | 46.5 | 15.6 | 830.75 |
| P33 | S28 | 2 | TB positive | Peru | Male | 179.3 | 135.2 | 3.9 | 8.9 | 3.99 | 1.145 | 131.4 | 5.6 | 0.25 | 1.09 | 0.95 | 4.38 | 45.1 | 176.55 | 26.65 | 27.45 | 804.9 |
| P34 | S23 | 3 | TB positive | Peru | Female | 227.1 | 87.3 | 8.2 | 3.45 | 3.925 | 0.315 | 61.55 | 1.55 | 0.62 | 1.525 | 4.355 | 7.27 | 37.75 | 207.65 | 8.25 | 36.95 | 365 |
| P35 | S34 | 3 | TB positive | Peru | Male | 288.85 | 178.95 | 10.4 | 10.4 | 16.435 | 0.49 | 184.05 | 3.45 | 0.01 | 2.785 | 0.01 | 10.25 | 87.2 | 117.05 | 29.9 | 18.7 | 739.05 |
| P36 | S25 | 3 | TB positive | Peru | Male | 251 | 206.25 | 4.1 | 2.9 | 10.565 | 0.425 | 123 | 2.9 | 0.01 | 4.285 | 0.01 | 8.43 | 73.5 | 99.55 | 37.9 | 15.85 | 338.95 |
| P37 | S27 | 2 | TB positive | Peru | Female | 233.7 | 88.15 | 3.65 | 9.1 | 4.365 | 1.03 | 238.3 | 3.15 | 0.41 | 2.595 | 0.95 | 6.63 | 96.75 | 168.8 | 25.3 | 26.95 | 834.1 |
| P38 | S26 | 3 | TB positive | Peru | Female | 296.05 | 105.15 | 8.45 | 1.8 | 6.28 | 0.53 | 191.65 | 2.75 | 0.505 | 2.52 | 1.145 | 5.055 | 100.35 | 116.65 | 38 | 19.35 | 140.05 |
| P39 | S22 | 3 | TB positive | Peru | Male | 299.1 | 135.55 | 15.35 | 3.35 | 9.945 | 0.385 | 338.15 | 3.65 | 0.01 | 2.93 | 0.01 | 6.65 | 328.4 | 159.85 | 155.95 | 35.65 | 4.8 |
| P40 | S37 | 3 | TB positive | Peru | Female | 248.8 | 55.9 | 3.25 | 0.9 | 5.07 | 0.365 | 73.6 | 1.55 | 0.08 | 0.78 | 0.315 | 2.4 | 21.2 | 70.4 | 9.4 | 10.7 | 28.8 |
| P41 | S51 | 0 | TB negative | Peru | Female | 268.55 | 71.7 | 1.4 | 0.35 | 2.24 | 0.37 | 54.3 | 0.75 | 0.32 | 1.71 | 1.095 | 5.08 | 32.7 | 50.5 | 11.8 | 9.9 | 7.25 |
| P42 | S50 | 0 | TB negative | Peru | Male | 258.4 | 53.8 | 2.75 | 0.7 | 6.905 | 0.305 | 165.7 | 1.7 | 0.34 | 1.105 | 0.97 | 3.215 | 55.1 | 82.7 | 20.2 | 14.7 | 11.35 |
| P43 | S52 | 0 | TB negative | Peru | Male | 317.9 | 110.6 | 2.15 | 0.5 | 4.28 | 0.24 | 73 | 1.8 | 0.01 | 2.94 | 0.01 | 7.77 | 33.35 | 97.4 | 22.55 | 16.3 | 15.45 |
| P44 | S45 | 0 | TB negative | Peru | Female | 418.15 | 173.9 | 11.95 | 1.15 | 20.625 | 0.545 | 116.15 | 3.4 | 0.01 | 4.265 | 0.01 | 12.195 | 52.75 | 124.7 | 27.25 | 20.6 | 3.95 |
| P45 | S44 | 0 | TB negative | Peru | Male | 234.25 | 105 | 12.7 | 1.15 | 23.905 | 0.7 | 211.85 | 4.3 | 0.01 | 3.07 | 0.01 | 13.285 | 82 | 214.1 | 37.7 | 31 | 3.4 |
| P46 | S42 | 0 | TB negative | Peru | Female | 267.2 | 100.3 | 5 | 0.85 | 9.72 | 0.385 | 92.8 | 2.3 | 0.01 | 2.215 | 0.01 | 6.87 | 41.65 | 112.75 | 16.05 | 15.3 | 8.1 |
| P47 | S43 | 0 | TB negative | Peru | Female | 209.7 | 56.1 | 3.75 | 0.5 | 9.26 | 0.4 | 96.8 | 1.95 | 0.12 | 1.155 | 1.075 | 6.66 | 52.4 | 149.3 | 25.7 | 25 | 7.75 |
| P48 | S41 | 0 | TB negative | Peru | Female | 360.95 | 163.4 | 11.7 | 1.4 | 16.295 | 0.395 | 136.6 | 2.4 | 0.01 | 3.71 | 0.01 | 9.475 | 65.2 | 132.15 | 25.6 | 21.15 | 9.8 |
| P49 | S46 | 0 | TB negative | Peru | Male | 269.1 | 60.6 | 9.4 | 1.15 | 27.585 | 0.58 | 163.5 | 2.95 | 0.625 | 1.405 | 2.23 | 4.955 | 47.05 | 130.65 | 15.95 | 19.45 | 6.05 |
| P50 | S49 | 0 | TB negative | Peru | Female | 274.2 | 98.85 | 10.7 | 0.65 | 13.82 | 0.28 | 50.05 | 1.95 | 0.01 | 3.02 | 0.01 | 13.86 | 18.1 | 138.65 | 11 | 22.15 | 4.9 |
| P51 | S48 | 0 | TB negative | Peru | Female | 450.1 | 152.4 | 16.25 | 1.7 | 27.39 | 0.64 | 112.85 | 3.45 | 0.325 | 2.56 | 2.665 | 16.165 | 43.25 | 253.75 | 18.6 | 49.55 | 11.05 |
| P52 | S47 | 0 | TB negative | Peru | Female | 230.45 | 44.65 | 4.2 | 0.65 | 6.3 | 0.58 | 181 | 2.3 | 0.6 | 2.495 | 1.575 | 5.215 | 68.85 | 118.95 | 24.85 | 19.05 | 5.55 |
| P53 | S54 | 0 | TB negative | Peru | Male | 325.55 | 64.4 | 9.85 | 1.7 | 15.565 | 0.525 | 137.55 | 2.4 | 0.25 | 1.46 | 0.245 | 4.095 | 50.5 | 94.75 | 22.4 | 17.05 | 33.25 |
| P54 | S55 | 0 | TB negative | Peru | Male | 400.55 | 93.15 | 10.7 | 4 | 32.265 | 0.5 | 137.65 | 2.15 | 0.78 | 1.26 | 3.37 | 5.605 | 55.05 | 131.8 | 15.15 | 19.25 | 234.15 |
| P55 | S70 | 3 | TB positive | Vietnam | Male | 415.1 | 176.95 | 5.9 | 1.15 | 15.47 | 0.27 | 184.05 | 3.85 | 0.035 | 2.65 | 0.01 | 11.04 | 38.15 | 110.25 | 16.5 | 15.05 | 5.15 |
| P56 | S60 | 0 | TB negative | Vietnam | Female | 346.4 | 137.2 | 9.6 | 1.45 | 14.475 | 0.29 | 118.3 | 2.35 | 0.01 | 3.13 | 0.01 | 10.76 | 83.05 | 164.1 | 32.9 | 24.1 | 12.2 |
| P57 | S61 | 0 | TB negative | Vietnam | Female | 543.6 | 98.6 | 23.25 | 3.85 | 53.93 | 0.825 | 226.25 | 3.55 | 1.915 | 3.165 | 8.715 | 13.62 | 100.05 | 298.25 | 27.55 | 35.9 | 11.2 |
| P58 | S72 | 0 | TB negative | Vietnam | Female | 464.9 | 196.1 | 9.5 | 1.4 | 27.7 | 0.55 | 130.35 | 2.7 | 0.01 | 2.12 | 0.01 | 9.895 | 50.35 | 102.4 | 16.8 | 15.6 | 7.65 |
| P59 | S69 | 2 | TB positive | Vietnam | Male | 292.6 | 82.65 | 2.05 | 0.9 | 7.54 | 0.245 | 161.8 | 0.95 | 0.01 | 1.505 | 0.055 | 4.175 | 60.55 | 84.45 | 24.3 | 13.75 | 13.7 |
| P60 | S76 | 3 | TB positive | Vietnam | Male | 323.35 | 130.85 | 7.85 | 1.6 | 18.84 | 0.57 | 146.6 | 3.2 | 0.01 | 2.945 | 0.01 | 7.175 | 79.2 | 143.75 | 31.55 | 23.2 | 3.15 |
| P61 | S66 | 3 | TB positive | Vietnam | Male | 419 | 62.25 | 7.35 | 2.2 | 30.025 | 0.5 | 248.3 | 2.2 | 0.635 | 3.53 | 0.88 | 11.885 | 105.6 | 166.5 | 43.1 | 24.35 | 10.25 |
| P62 | S65 | 2 | TB positive | Vietnam | Female | 304.75 | 100.2 | 9.9 | 1.35 | 21.79 | 0.285 | 95.25 | 1.85 | 0.01 | 4.03 | 0.01 | 15.59 | 34.15 | 161.2 | 17.1 | 25 | 24.95 |
| P63 | S67 | 3 | TB positive | Vietnam | Male | 256 | 81.75 | 7.95 | 0.95 | 16.68 | 0.33 | 134.9 | 2.75 | 0.01 | 4.3 | 0.01 | 10.855 | 80.55 | 138.15 | 40.5 | 20.55 | 3.75 |
| P64 | S64 | 2 | TB positive | Vietnam | Male | 385.3 | 75.05 | 10.75 | 1.75 | 15.575 | 0.55 | 168.15 | 1.85 | 0.775 | 1.765 | 1.705 | 5.94 | 63.1 | 128.5 | 16.05 | 18.65 | 8.8 |
| P65 | S71 | 0 | TB negative | Vietnam | Male | 563.1 | 206.45 | 14.8 | 2.85 | 33.595 | 0.765 | 199.9 | 5.2 | 0.01 | 5.215 | 0.01 | 24.505 | 94.9 | 191.65 | 40.35 | 28.85 | 12.9 |
| P66 | S62 | 2 | TB positive | Vietnam | Female | 353.65 | 99.45 | 7.9 | 2.8 | 28.14 | 0.315 | 166.5 | 1.3 | 1.375 | 2.02 | 2.96 | 3.83 | 61.2 | 91.8 | 20.55 | 15.25 | 18.3 |
| P67 | S59 | 2 | TB positive | Vietnam | Male | 382.25 | 110.9 | 5.1 | 1.05 | 16.31 | 0.305 | 198.2 | 2.45 | 0.01 | 3.8 | 0.01 | 8.91 | 125.85 | 153 | 56.15 | 22.85 | 4.2 |
| P68 | S73 | 2 | TB positive | Vietnam | Female | 352.65 | 127.35 | 7.9 | 1.05 | 11.14 | 0.355 | 92.95 | 1.45 | 0.01 | 3.43 | 0.01 | 10.22 | 49.95 | 83.5 | 18.75 | 13.4 | 4.55 |
| P69 | S74 | 3 | TB positive | Vietnam | Male | 391.55 | 85.4 | 7.25 | 1.4 | 17.745 | 0.33 | 121.1 | 1.95 | 0.155 | 2.295 | 0.545 | 8.945 | 76.45 | 129.7 | 24.6 | 18.5 | 7.5 |
| P70 | S68 | 3 | TB positive | Vietnam | Male | 345.05 | 45.5 | 4.75 | 0.95 | 10.78 | 0.34 | 147.9 | 1.65 | 0.52 | 3.605 | 1.24 | 9.365 | 77.2 | 148.1 | 38.1 | 23.1 | 5.45 |
| P71 | S53 | 3 | TB positive | Vietnam | Female | 353.25 | 62.65 | 5.35 | 0.95 | 10.77 | 0.34 | 109.65 | 0.95 | 1 | 1.69 | 3.88 | 5.125 | 50.9 | 122.15 | 14.45 | 18.7 | 6.5 |
| P72 | S57 | 3 | TB positive | Vietnam | Male | 387.75 | 57.45 | 6.9 | 1.05 | 27.145 | 0.37 | 235.15 | 2.05 | 0.73 | 1.61 | 2.375 | 5.205 | 95.45 | 119.05 | 34.6 | 18.1 | 7.8 |
| P73 | S58 | 3 | TB positive | Vietnam | Male | 379.75 | 58.3 | 14.7 | 2.15 | 46.595 | 0.58 | 300.8 | 2.05 | 1.065 | 1.505 | 4.7 | 5.85 | 107.5 | 187.75 | 38.7 | 28.05 | 5 |
| P74 | S56 | 2 | TB positive | Vietnam | Male | 440.55 | 99.75 | 8.95 | 1.6 | 22.39 | 0.46 | 158.4 | 2.4 | 0.01 | 3.995 | 0.01 | 12.495 | 77.6 | 120.9 | 34 | 18.05 | 11.4 |
| P75 | S77 | 2 | TB positive | Vietnam | Male | 296.05 | 57.75 | 8.35 | 0.95 | 14.68 | 0.345 | 186.9 | 1.65 | 0.36 | 2.45 | 0.785 | 6.595 | 84.9 | 96.2 | 33.85 | 16.15 | 4.85 |
| P76 | S78 | 3 | TB positive | Vietnam | Female | 340.75 | 92.5 | 10.05 | 1.2 | 10.375 | 0.335 | 102.3 | 0.9 | 0.645 | 3.41 | 1.19 | 9.48 | 55.3 | 109.3 | 19.95 | 21.5 | 6.15 |
| P77 | S63 | 2 | TB positive | Vietnam | Male | 410.1 | 105.1 | 8.7 | 1.9 | 30.585 | 0.665 | 277.1 | 2.35 | 0.98 | 2.355 | 4.045 | 7.785 | 115.6 | 208.8 | 36 | 29.25 | 4.4 |
| P78 | S75 | 2 | TB positive | Vietnam | Male | 482.3 | 214.95 | 14.35 | 3.2 | 31.605 | 0.615 | 120.65 | 2.6 | 1 | 3.82 | 3.6 | 19.79 | 67.6 | 356.85 | 21.7 | 51.5 | 39.35 |
| P79 | S80 | 0 | TB negative | Peru | Male | 202.15 | 96.75 | 4.55 | 2.3 | 11.06 | 0.4 | 218.45 | 2.7 | 0.055 | 1.405 | 0.01 | 4.48 | 84.3 | 103.05 | 35.8 | 16.7 | 131.4 |
| P80 | S79 | 0 | TB negative | Peru | Male | 190.25 | 50.35 | 2.05 | 1.65 | 7.62 | 0.505 | 148.15 | 2.2 | 0.205 | 0.825 | 0.475 | 3.065 | 51.65 | 72.65 | 20.75 | 11.2 | 49.2 |

| ID | Patient_ID | Time | Age | Cavitary | Sex | Total cholesterol (μg/mL) | Free cholesterol (μg/mL) | Esterified cholesterol (μg/mL) | Cholest-4-en-3-one (ng/mL) | 27-hydroxycholesterol (ng/mL) | 27-hydroxycholest-4-en-3-one (ng/mL) | 3β-hydroxycholest-5-enoic acid (ng/mL) | 3-oxocholest-4-enoic acid (ng/mL) |
|---|---|---|---|---|---|---|---|---|---|---|---|---|---|
| 0207A Pretreatment | 1 | Before | 35 | Yes | Female | 83.13 | 14.25 | 68.88 | 3200.00 | 477.00 | 271.00 | 2720.00 | 291.00 |
| 0633C Pretreatment | 2 | Before | 28 | No | Female | 30.96 | 4.14 | 26.82 | 257.00 | 32.20 | 13.40 | 170.00 | 7.47 |
| 0005A Pretreatment | 3 | Before | 30 | Yes | Female | 23.09 | 3.33 | 19.76 | 357.00 | 105.00 | 86.10 | 534.00 | 62.00 |
| 0006A Pretreatment | 4 | Before | 27 | Yes | Female | 32.91 | 7.45 | 25.46 | 154.00 | 136.00 | 61.90 | 939.00 | 26.60 |
| 0134A Pretreatment | 5 | Before | 24 | Yes | Female | 96.56 | 18.21 | 78.34 | 82.00 | 67.80 | 0.10 | 322.00 | 1.81 |
| 0197A Pretreatment | 6 | Before | 22 | Yes | Female | 22.67 | 3.24 | 19.42 | 274.00 | 87.50 | 14.50 | 279.00 | 9.59 |
| 0071A Pretreatment | 7 | Before | 20 | No | Female | 58.06 | 5.75 | 52.31 | 53.80 | 0.10 | 0.10 | 0.10 | 0.10 |
| 0031A Pretreatment | 8 | Before | 22 | Yes | Female | 16.02 | 4.36 | 11.66 | 169.00 | 54.40 | 0.10 | 211.00 | 0.24 |
| 0039A Pretreatment | 9 | Before | 23 | Yes | Female | 30.91 | 4.63 | 26.28 | 73.10 | 22.50 | 6.21 | 133.00 | 2.39 |
| 0023A Pretreatment | 10 | Before | 25 | Yes | Female | 54.34 | 9.25 | 45.09 | 213.00 | 145.00 | 79.30 | 908.00 | 45.10 |
| 0183A Pretreatment | 11 | Before | 19 | Yes | Female | 27.26 | 1.52 | 25.74 | 33.30 | 28.00 | 29.90 | 326.00 | 7.41 |
| 0067A Pretreatment | 12 | Before | 22 | Yes | Female | 53.88 | 8.09 | 45.79 | 250.00 | 1880.00 | 165.00 | 11000.00 | 61.90 |
| 0138A Pretreatment | 13 | Before | 32 | No | Female | 15.13 | 1.44 | 13.68 | 44.20 | 3.19 | 0.10 | 95.60 | 3.72 |
| 0068A Pretreatment | 14 | Before | 27 | Yes | Female | 75.94 | 24.07 | 51.87 | 1850.00 | 98.50 | 378.00 | 1710.00 | 242.00 |
| 0020A Pretreatment | 15 | Before | 19 | Yes | Female | 32.02 | 7.13 | 24.90 | 492.00 | 47.10 | 157.00 | 579.00 | 104.00 |
| 0627C Pretreatment | 16 | Before | 41 | No | Male | 90.65 | 12.17 | 78.48 | 1070.00 | 261.00 | 114.00 | 937.00 | 39.40 |
| 0037A Pretreatment | 17 | Before | 35 | Yes | Male | 104.77 | 12.03 | 92.74 | 2630.00 | 228.00 | 168.00 | 1340.00 | 161.00 |
| 0603C Pretreatment | 18 | Before | 25 | Yes | Male | 19.66 | 7.18 | 12.48 | 132.00 | 48.60 | 0.10 | 1360.00 | 1.88 |
| 0264A Pretreatment | 19 | Before | 43 | Yes | Male | 25.54 | 3.25 | 22.29 | 282.00 | 42.30 | 23.70 | 376.00 | 15.50 |
| 0639C Pretreatment | 20 | Before | 29 | No | Male | 47.90 | 9.90 | 38.00 | 528.00 | 539.00 | 18.10 | 1060.00 | 5.85 |
| 0010A Pretreatment | 21 | Before | 22 | Yes | Male | 36.30 | 5.84 | 30.47 | 490.00 | 55.10 | 65.90 | 452.00 | 45.70 |
| 0635C Pretreatment | 22 | Before | 33 | Yes | Male | 7.85 | 3.50 | 4.35 | 933.00 | 36.30 | 39.90 | 298.00 | 57.50 |
| 0008A Pretreatment | 23 | Before | 22 | Yes | Male | 13.06 | 2.29 | 10.78 | 170.00 | 27.40 | 24.50 | 309.00 | 29.40 |
| 0607C Pretreatment | 24 | Before | 42 | Yes | Male | 88.71 | 3.08 | 85.63 | 177.00 | 28.00 | 7.12 | 93.10 | 1.59 |
| 0065A Pretreatment | 25 | Before | 34 | Yes | Male | 23.41 | 6.31 | 17.09 | 408.00 | 299.00 | 56.90 | 1680.00 | 34.50 |
| 0156A Pretreatment | 26 | Before | 23 | Yes | Male | 18.77 | 5.54 | 13.22 | 545.00 | 140.00 | 46.70 | 566.00 | 19.10 |
| 0174A Pretreatment | 27 | Before | 31 | Yes | Male | 11.31 | 0.97 | 10.34 | 60.00 | 9.79 | 0.10 | 2440.00 | 0.84 |
| 0920A Pretreatment | 28 | Before | 22 | Yes | Male | 39.30 | 6.52 | 32.79 | 574.00 | 37.50 | 71.40 | 512.00 | 30.80 |
| 0915A Pretreatment | 29 | Before | 28 | No | Male | 58.33 | 7.40 | 50.93 | 67.70 | 2.31 | 0.10 | 45.70 | 0.35 |
| 0042A Pretreatment | 30 | Before | 39 | Yes | Male | 49.12 | 6.42 | 42.70 | 191.00 | 115.00 | 13.10 | 1120.00 | 5.40 |
| 0207A 8 weeks of treatm | 1 | After | 35 | Yes | Female | 41.98 | 2.61 | 39.37 | 34.40 | 9.16 | 0.10 | 46.90 | 0.10 |
| 0633C 8 weeks of treatm | 2 | After | 28 | No | Female | 10.52 | 5.43 | 5.09 | 45.20 | 10.10 | 0.10 | 59.60 | 0.10 |
| 0005A 8 weeks of treatm | 3 | After | 30 | Yes | Female | 19.91 | 3.59 | 16.32 | 94.00 | 118.00 | 0.10 | 704.00 | 0.37 |
| 0006A 8 weeks of treatm | 4 | After | 27 | Yes | Female | 16.06 | 5.17 | 10.89 | 40.00 | 120.00 | 0.10 | 374.00 | 0.10 |
| 0134A 8 weeks of treatm | 5 | After | 24 | Yes | Female | 56.01 | 5.46 | 50.55 | 49.40 | 19.90 | 0.10 | 61.50 | 0.10 |
| 0197A 8 weeks of treatm | 6 | After | 22 | Yes | Female | 14.42 | 2.92 | 11.50 | 134.00 | 74.50 | 0.10 | 293.00 | 0.10 |
| 0071A 8 weeks of treatm | 7 | After | 20 | No | Female | 14.20 | 1.64 | 12.56 | 56.00 | 0.10 | 0.10 | 0.10 | 0.10 |
| 0031A 8 weeks of treatm | 8 | After | 22 | Yes | Female | 20.80 | 2.50 | 18.30 | 63.20 | 71.30 | 0.10 | 155.00 | 0.10 |
| 0039A 8 weeks of treatm | 9 | After | 23 | Yes | Female | 29.32 | 4.29 | 25.03 | 17.20 | 105.00 | 0.10 | 439.00 | 0.10 |
| 0023A 8 weeks of treatm | 10 | After | 25 | Yes | Female | 33.19 | 5.41 | 27.78 | 64.70 | 155.00 | 0.10 | 594.00 | 0.10 |
| 0183A 8 weeks of treatm | 11 | After | 19 | Yes | Female | 17.56 | 3.58 | 13.98 | 65.20 | 62.70 | 0.10 | 194.00 | 0.04 |
| 0067A 8 weeks of treatm | 12 | After | 22 | Yes | Female | 35.30 | 8.17 | 27.13 | 150.00 | 161.00 | 0.10 | 484.00 | 0.13 |
| 0138A 8 weeks of treatm | 13 | After | 32 | No | Female | 23.90 | 2.04 | 21.87 | 32.90 | 34.10 | 0.10 | 175.00 | 0.10 |
| 0068A 8 weeks of treatm | 14 | After | 27 | Yes | Female | 77.52 | 7.66 | 69.85 | 199.00 | 50.70 | 0.10 | 265.00 | 0.17 |
| 0020A 8 weeks of treatm | 15 | After | 19 | Yes | Female | 55.79 | 6.20 | 49.59 | 52.90 | 97.10 | 0.10 | 232.00 | 0.10 |
| 0627C 8 weeks of treatm | 16 | After | 41 | No | Male | 16.53 | 2.14 | 14.39 | 74.90 | 83.20 | 0.10 | 186.00 | 0.10 |
| 0037A 8 weeks of treatm | 17 | After | 35 | Yes | Male | 30.19 | 4.04 | 26.16 | 65.90 | 55.40 | 0.10 | 135.00 | 0.10 |
| 0603C 8 weeks of treatm | 18 | After | 25 | Yes | Male | 65.67 | 8.32 | 57.35 | 211.00 | 203.00 | 0.10 | 5620.00 | 2.09 |
| 0264A 8 weeks of treatm | 19 | After | 43 | Yes | Male | 40.20 | 4.58 | 35.62 | 46.10 | 14.60 | 0.10 | 859.00 | 0.41 |
| 0639C 8 weeks of treatm | 20 | After | 29 | No | Male | 38.31 | 4.70 | 33.61 | 53.10 | 3.07 | 0.10 | 18.30 | 0.21 |
| 0010A 8 weeks of treatm | 21 | After | 22 | Yes | Male | 12.89 | 4.06 | 8.82 | 72.00 | 72.60 | 0.10 | 467.00 | 0.10 |
| 0635C 8 weeks of treatm | 22 | After | 33 | Yes | Male | 14.82 | 6.49 | 8.33 | 59.10 | 41.90 | 0.10 | 184.00 | 0.10 |
| 0008A 8 weeks of treatm | 23 | After | 22 | Yes | Male | 23.16 | 6.27 | 16.89 | 85.10 | 199.00 | 0.10 | 2200.00 | 1.26 |
| 0607C 8 weeks of treatm | 24 | After | 42 | Yes | Male | 7.98 | 2.57 | 5.41 | 35.40 | 9.91 | 0.10 | 69.80 | 0.10 |
| 0065A 8 weeks of treatm | 25 | After | 34 | Yes | Male | 44.97 | 7.86 | 37.11 | 125.00 | 506.00 | 0.10 | 7810.00 | 3.02 |
| 0156A 8 weeks of treatm | 26 | After | 23 | Yes | Male | 15.96 | 1.74 | 14.21 | 27.40 | 3.30 | 0.10 | 0.10 | 0.10 |
| 0174A 8 weeks of treatm | 27 | After | 31 | Yes | Male | 14.60 | 2.17 | 12.43 | 33.10 | 1.42 | 0.10 | 0.10 | 0.10 |
| 0920A 8 weeks of treatm | 28 | After | 22 | Yes | Male | 18.02 | 4.21 | 13.80 | 54.40 | 32.30 | 0.10 | 335.00 | 0.10 |
| 0915A 8 weeks of treatm | 29 | After | 28 | No | Male | 16.12 | 1.87 | 14.25 | 42.60 | 14.40 | 0.10 | 53.70 | 0.10 |
| 0042A 8 weeks of treatm | 30 | After | 39 | Yes | Male | 11.40 | 3.06 | 8.34 | 91.70 | 31.60 | 0.10 | 348.00 | 0.10 |
